# Co-ordinated Ras and Rac activity shapes macropinocytic cups and enables phagocytosis of geometrically diverse bacteria

**DOI:** 10.1101/763748

**Authors:** Catherine M. Buckley, Henderikus Pots, Aurelie Gueho, Ben A. Phillips, Bernd Gilsbach, David Traynor, Anton Nikolaev, Thierry Soldati, Andrew J. Parnell, Arjan Kortholt, Jason S. King

## Abstract

Engulfment of extracellular material by phagocytosis or macropinocytosis depends on the ability of cells to generate specialised cup shaped protrusions. To effectively capture and internalise their targets, these cups are organised into a ring or ruffle of actin-driven protrusion encircling a non-protrusive interior domain. These functional domains depend on the combined activities of multiple Ras and Rho family small GTPases, but how their activities are integrated and differentially regulated over space and time is unknown. Here, we show that the amoeba *Dictyostelium discoideum* coordinates Ras and Rac activity using the multidomain protein RGBARG (<u>R</u>CC1, Rho<u>G</u>EF, <u>BAR</u> and Ras<u>G</u>AP-containing protein). We find RGBARG uses a tripartite mechanism of Ras, Rac and phospholipid interactions to localise at the protruding edge and interface with the interior of both macropinocytic and phagocytic cups. There, RGBARG shapes the protrusion by driving Rac activation at the rim whilst suppressing expansion of the active Ras interior domain. Consequently, cells lacking RGBARG form enlarged, flat interior domains unable to generate large macropinosomes. During phagocytosis, we find that disruption of *RGBARG* causes a geometry-specific defect in engulfing rod-shaped bacteria and ellipsoidal beads. This demonstrates the importance of co-ordinating small GTPase activities during engulfment of more complex shapes and thus the full physiological range of microbes, and how this is achieved in a model professional phagocyte.

## Introduction

The capture and engulfment of extracellular material serves a number of important cellular functions. Best understood is the clearance of pathogenic microbes or apoptotic cells by phagocytic immune cells, but the engulfment of fluid by the related process of macropinocytosis also plays important functions by allowing cells to capture antigens or other factors from their environment such as nutrients to support growth (Bloomfield and Kay, 2016; Commisso et al., 2013; Norbury et al., 1995; Sallusto et al., 1995; Swanson and King, 2019).

To capture extracellular material, cells must encircle and isolate their target within a vesicle. This can be achieved by several mechanisms, but the best understood and evolutionarily widespread involves the extension of a circular cup or ruffle-shaped protrusion from the cell surface to enwrap and internalize the target (Buckley and King, 2017; Kaplan, 1977; Swanson, 2008; Veltman et al., 2016). Whilst many components of cup formation have been identified, how they are co-ordinated in space and time is poorly understood. Here we describe a novel mechanism used by the amoebae *Dictyostelium discoideum* to integrate different signaling elements and form complex cup-shaped protrusions that efficiently mediate engulfment.

Macropinocytic and phagocytic protrusions are formed by localised actin polymerisation at the plasma membrane, using much of the same machinery that generates pseudopods and lamellipodia during cell migration (King and Kay, 2019; Swanson, 2008). Whilst migratory protrusions only need the cell to define a simple patch of actin polymerisation, forming a cup requires a higher level of organisation, with the protrusive activity restricted to a ring encircling a static interior domain. During phagocytosis this is aided by the presence of a particle to act as a physical scaffold and locally activate receptors. These interactions are proposed to guide engulfment by a zippering mechanism (Griffin et al., 1975; Tollis et al., 2010). However, macropinocytic cups self-organise with an almost identical structure in the absence of any external spatial cues (Veltman et al., 2016). Cup formation can therefore occur spontaneously by the intrinsic dynamics of the underlying signaling.

Recent studies in *Dictyostelium* proposed a model whereby the cup interior is defined by spontaneous localised activation of the small GTPase Ras and consequent accumulation of the phospholipid PIP_3_ (Veltman et al., 2016). This patch appears to restrict actin polymerisation to its periphery to create a protrusive ring. How this is achieved is unknown, but in at least *Dictyostelium* it may depend on the activity of the PIP_3_-activated Protein kinase B/Akt (Williams et al., 2019). Both active Ras and PIP_3_ also accumulate at cups in mammalian cells (Araki et al., 2007; Marshall et al., 2001; Vieira et al., 2001) and Ras activation is sufficient to drive ruffling and macropinocytosis in cancer cells (Bar-Sagi and Feramisco, 1986; Commisso et al., 2013). PI3K inhibition also completely blocks macropinocytosis (Amyere et al., 2000; Araki et al., 1996; Hoeller et al., 2013; Veltman et al., 2014) as well as phagocytosis of large particles by macrophages (Araki et al., 1996; Cox et al., 1999; Schlam et al., 2015). Ras and PIP_3_ therefore play a general role in macropinosome and phagosome organisation across evolution.

Other small GTPases are also involved in cup formation. Active Rac1 overlaps with Ras activity in the cup interior in both macrophages and *Dictyostelium* (Hoppe and Swanson, 2004; Veltman et al., 2016). Rac1 is a direct activator of the SCAR/WAVE complex, which drives activation of actin polymerisation via the ARP2/3 complex (Eden et al., 2002; Machesky and Insall, 1998). Consistent with this, Rac1 is required for macropinosome formation in dendritic cells (West et al., 2000) and optogenetic Rac1 activation is sufficient to drive ruffling and macropinocytosis in macrophages (Fujii et al., 2013). Expression of constitutively active Rac1 also leads to excessive actin at macropinocytic cups in *Dictyostelium* (Dumontier et al., 2000). Therefore, whilst Ras appears to define the cup interior, Rac1 is important for regulating actin protrusions, as it is does during cell migration.

The presence of active Rac1 throughout the cup interior is at odds with the tightly restricted SCAR/WAVE activity and protrusion at the extending rim (Veltman et al., 2016). A further layer of regulation must therefore exist. This is likely provided by the small GTPase CDC42 which is also required for Fc-γ-receptor mediated phagocytosis and collaborates with Rac1 during engulfment of large particles (Caron and Hall, 1998; Cox et al., 1997; Massol et al., 1998; Schlam et al., 2015). In contrast to Rac1, active CDC42 is restricted to the protrusive cup rim in macrophages indicating differential regulation and functionality (Hoppe and Swanson, 2004). In *Dictyostelium* however, no clear CDC42 orthologue has been identified.

Cup formation requires integrated spatio-temporal control over multiple GTPases. This must be able to self-organise in the absence of external cues during macropinocytosis, and robust enough to phagocytose physiological targets of varying size and shape. Small GTPase activity is controlled by a large family of proteins such as GTPase Exchange Factors (GEFs) which promote the GTP-bound active form, and GTPase Activating Proteins (GAPs) which stimulate hydrolysis and transition to a GDP-bound inactive state. In this study, we characterise a previously unstudied dual GEF and GAP protein in *Dictyostelium* that integrates Ras, Rac and lipid signaling. This provides a mechanism to coordinate the cup interior with the protrusive rim, allowing efficient macropinosome formation and the engulfment of diverse bacteria of differing geometry.

## Results

### Identification of a novel BAR-domain containing protein recruited to cups

Our initial hypothesis was that cells may use the different membrane curvature at the protrusive rim compared to the cup base to recognise and differentially regulate cup shape. Membrane curvature can recruit specific proteins containing BAR (Bin-Amphiphysin-Rvs) domains (Peter et al., 2004). These are often found in multidomain proteins, including several involved in GTPase regulation and trafficking (Aspenstrom, 2014). To identify candidate proteins involved in macropinocytosis, we therefore searched the *Dictyostelium* genome for BAR domain-containing proteins. Excluding proteins of known localisation or function, we systematically cloned each candidate and expressed them as both N- and C-terminal GFP-fusions in axenic Ax2 cells. Using this strategy we successfully cloned 9 previously uncharacterised BAR-containing proteins and observed their localisation in live cells by fluorescence microscopy. Of these, 6 were expressed at detectable levels (Figure 1A).

**Figure 1:**
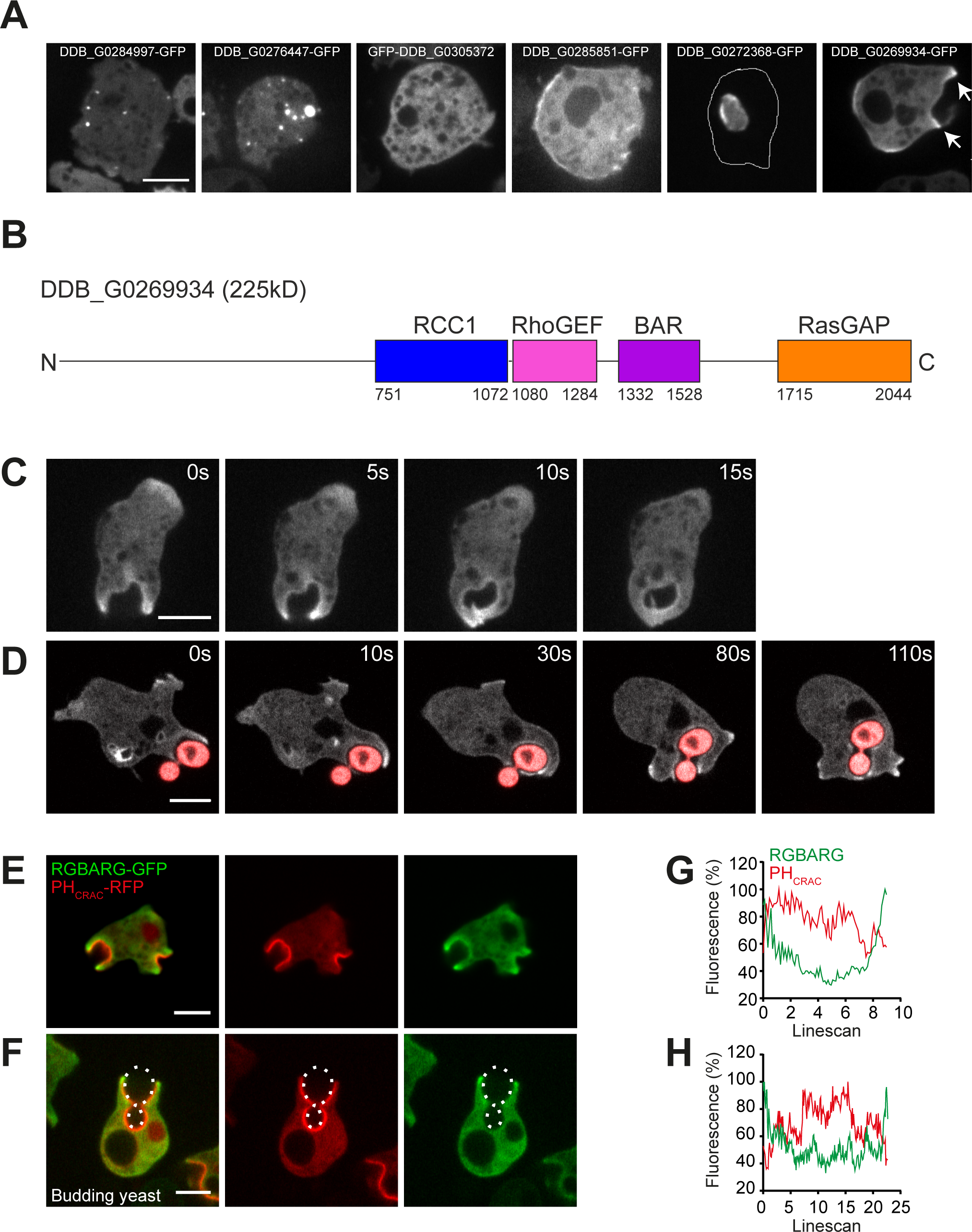
Identification of BAR domain proteins associated with macropinocytosis. (A) Uncharac-terised BAR-domain containing proteins were expressed as GFP-fusions in Ax2 Dictyostelium cells. Images are maximum intensity projections of confocal Z-stacks. (B) Illustrates the domain organisation of DDB_G0269934/RGBARG. (C) Time series of spinning disc images of RGBARG-GFP recruitment during macropinocytosis and (D) phagocytosis of TRITC-labeled heat-killed budding yeast. (E) and (D) Show the localization of RGBARG-GFP relative to the PIP3 reporter PHCRAC-RFP during macropino-cytosis and phagocytosis respectively. (G) and (H) show the relative intensity profiles of each fluores-cent protein in these images. Linescans were drawn along the cup interior from protrusive tip-to-tip. All scale bars denote 5 μm.

DDB_G0284997, DDB_G0305372 and DDB_G0285851 were associated with small puncta at the plasma membrane, consistent with the well-characterised role of BAR domain proteins in clathrin mediated endocytosis (Dawson et al., 2006). DDB_G0276447 localised to intracellular vesicles too small to be macropinosomes, and GFP-DDB_G0272368 was exclusively observed in the nucleus. Only one of the proteins tested (DDB_G0269934) localised to what appeared to be the protrusive regions of macropinocytic cups.

DDB_G0269934 is a 223 kDa multidomain protein and also contains Regulator of Chromatin Condensation (RCC1), RhoGEF and RasGAP domains (Figure 1B). DDB_G0269934 has not previously been described and due to its domain organisation we will subsequently refer to it as RGBARG (<u>R</u>CC1, <u>G</u>EF, <u>BAR</u> and <u>G</u>AP domain containing protein). How Ras and Rac activity are coordinated in space and time to generate a 3-dimensional cup shaped protrusion is not known. Combining BAR, GEF and GAP activities in a single protein potentially provides an elegant mechanism to organise engulfment. Therefore the function and regulation of RGBARG was investigated in detail.

Examining RGBARG-GFP dynamics by timelapse fluorescence microscopy confirmed strong enrichment at the protrusive rim of both macropinocytic and phagocytic cups that disappeared rapidly after engulfment (Figure 1C and D, Videos 1 and 2). Co-expression with the PIP_3_ reporter PH_CRAC_-RFP that demarks the cup interior confirmed RGBARG-GFP localised specifically to the periphery of this signaling domain (Figure 1E-H, and Video 3). Importantly, this differs from the RasGAP NF1, which localises throughout the cup interior (Bloomfield et al., 2015). RGBARG may therefore play a specific role in organising cup dynamics and engulfment.

### RGBAR regulates cup signaling and macropinosome formation

To test for a functional role in engulfment a 3.6 Kb central section of *RGBARG* gene (containing the RCC1 RhoGEF and BAR domains) was deleted and replaced with a blasticidin selection cassette by homologous recombination (Figure 2 supplement 1). Independent clones were isolated (JSK02 and 03) and shared comparable phenotypes. In the following experiments strain JSK02 was used unless otherwise stated with effects of *RGBARG* mutation validated by rescue experiments.

To check for defects in macropinosome formation, cells were incubated with FITC-dextran, a pH sensitive dye that is quenched at low pH. As macropinosomes acidify in under two minutes in *Dictyostelium*, and images were acquired within the 30 minutes required to transit to neutral post-lysosomes, intracellular FITC-dextran is only visible in nascent macropinosomes (Figure 2A). From confocal Z-stacks of live cells, we found that *RGBARG*^*-*^ cells formed significantly smaller macropinosomes than parental controls, measuring 0.5±0.1 μm^3^ compared to 1.5±0.2 μm^3^ in Ax2 (Figure 2B). This phenotype could be completely rescued by re-expression of RGBARG-GFP. *RGBARG*^*-*^ cells also produced more macropinosomes (2.6±0.2 per cell compared to 2.2 ±0.1 for Ax2) leading to no net change in either total fluid uptake or axenic growth (Figure 2C-E). RGBARG is therefore functionally important for the dynamics of macropinocytosis but not essential for engulfment.

**Figure 2:**
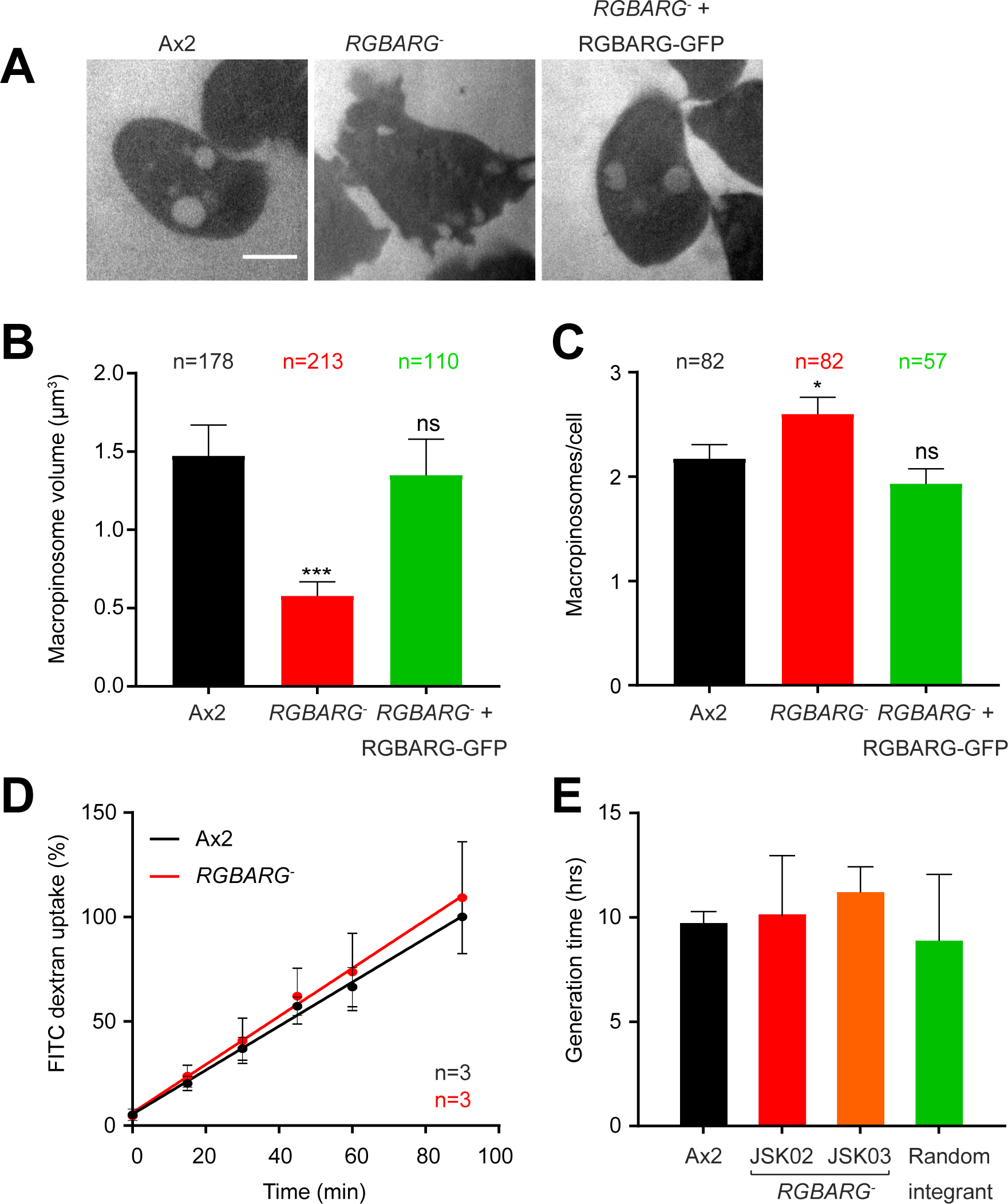
RGBARG-cells produce more, but smaller macropinosomes. (A) Confocal images of the indicated cell lines incubated in FITC-dextran for 10 minutes. The pH-sensitive FITC is only visible in pre-acidification macropinosomes <2 minutes after formation. The average volume of macropinosomes formed is shown in (B) and the number of macropinosomes per cell is shown in (C). n indicates the total number of macropinosomes or cells measured over 3 independent experiments. (D) Total fluid uptake measured by FITC dextran uptake over time, measured in a fluorimeter. (E) Growth rate of RGBARG mutants in axenic culture compared to the Ax2 parental cell line and a random integrant control from 3 independent experiments. All graphs show means ± SEM, * P<0.05, *** P<0.001 as determined by Students T-test.macropinosomes

To understand why *RGBARG*^*-*^ cells form smaller macropinosomes we followed their formation by fluorescence microscopy. Using the PH_CRAC_-GFP reporter we found in *RGBARG-*cells had larger and more numerous patches of PIP_3_ than controls, averaging 3.8±1.1 patches per confocal section with an average length of 5.5±2.4 μm compared to 1.6±0.6 patches averaging 4.4±1.2 μm in Ax2 (Figure 3A-C). Similarly enlarged patches were also observed using the active Ras reporter GFP-RBD (the Ras binding domain of Raf1, Figure 3B-C and Figure 3 Supplement 1A). During these experiments we also noted that *RGBARG*^*-*^ cells a mild cytokinesis defect with 10 ± % containing >2 nuclei, compared to 5% of controls. This is consistent with the cytokinesis defects described in *PTEN* mutants which also have excessive PIP_3_ accumulation (Janetopoulos et al., 2005) and all multinucleate cells were excluded from analysis.

**Figure 3:**
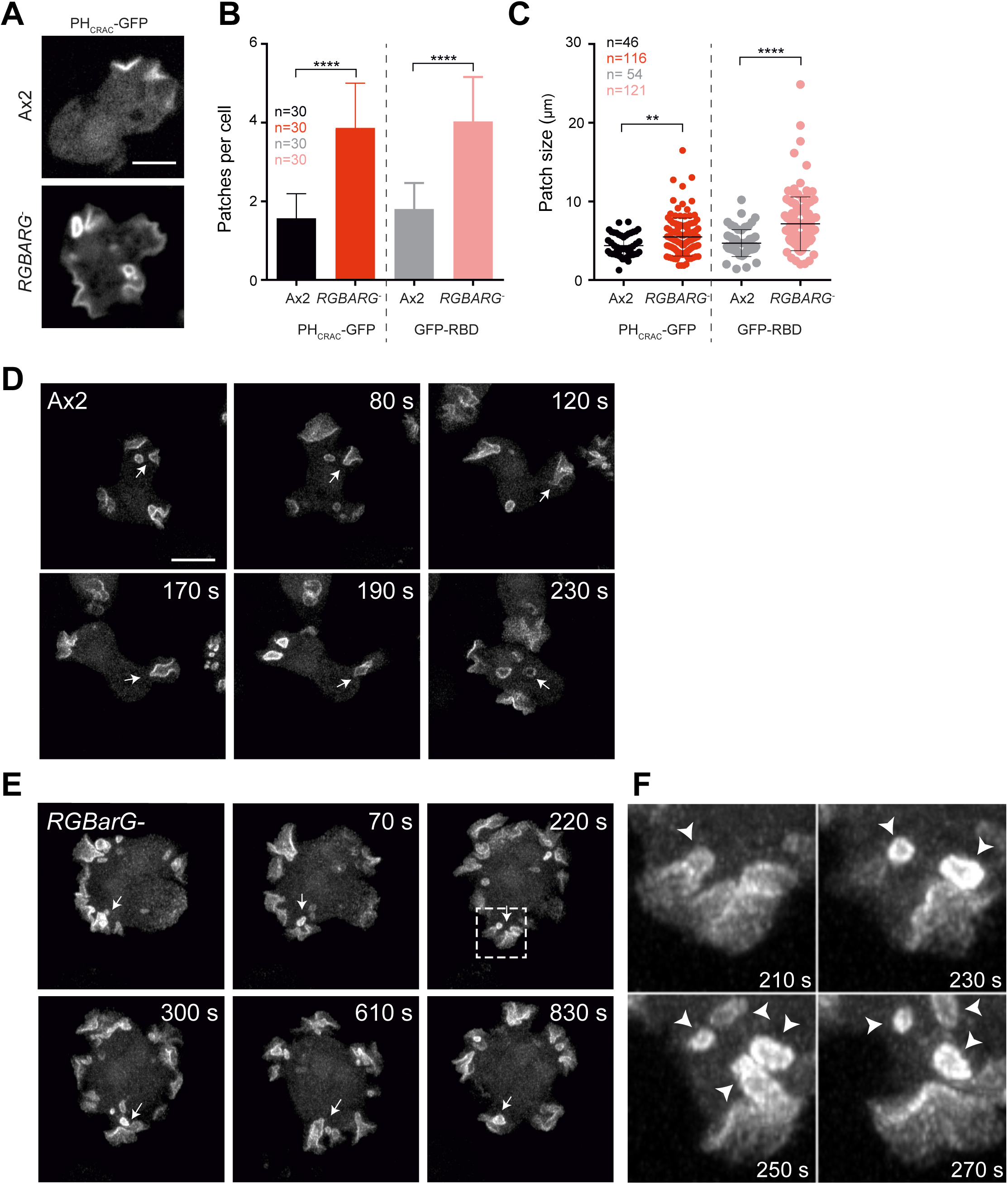
Dynamics of macropinosome formation in RGBARG-cells. (A) Membrane localization of the PIP3 probe PHCRAC-GFP. Images are single confocal sections taken by spinning disc microscopy. The number of patches of PHCRAC-GFP or the active Ras probe GFP-RBD per cell in single plane through the middle of each cell is quantified in (B). The average size of each patch is shown in (C). n is the total number of cells or patches measured over 3 independent experiments. Error bars show the mean ± standard deviation, ** P<0.01, *** P<0.001, Mann-Whitney T-test. (D) and (E) Time series of a maximum intensity projection through the entire depth of PHCRAC-GFP expressing cells. (D) Shows Ax2 cells, indicating the formation, closure and subsequent extinction of PHCRAC-GFP patch at the cell surface (arrow). (E) Shows an equivalent movie of RGBARG-cells, where the PHCRAC-GFP patch remains after closure (arrow). (F) Is an enlargement of the boxed area in (E), showing multiple vesicles forming from a single, large PHCRAC-GFP patch (arrowheads). All scale bars indicate 5 μm.macropinosomes

To understand how the enlarged Ras and PIP_3_ patches in *RGBARG*^-^ cells give rise to smaller macropinosomes we studied their formation over time in 3D. As recently described, the macropinocytic cups of Ax2 cells form by expanding around a defined spontaneous patch of PIP_3_ (Veltman et al., 2016). These cups subsequently close, usually forming one, or sometimes two, large macropinosomes accompanied by termination of PIP_3_ signaling on both the new vesicle and the cell surface (Figure 3D and Video 4). This process is relatively consistent, with each PIP_3_ patch lasting an average of 150 seconds (Figure 3 Supplement 1B). In *RGBARG*^-^ cells however, whilst PH_CRAC_-GFP still disappeared from internalised vesicles, the plasma membrane domains were much more stable. Whilst PIP_3_ patches frequently split, they rarely dissipated completely and often lasted longer than each of the 30 minute movies recorded (Figure 3E and Video 5). It was therefore not possible to meaningfully measure the lifetime of surface PIP_3_ (and by extension Ras) signaling in *RGBARG*^-^ cells. As RGBARG is restricted to the periphery of Ras signaling domains it appears to restrict both lateral expansion of activated Ras and termination of Ras/PIP_3_ signaling upon cup completion.

Although extinction of PIP_3_ signaling did not accompany cup closure in *RGBARG*^-^ cells, numerous small vesicles could be observed continuously budding from the base of the ruffles when folds of membrane collapsed in on themselves. This explains why these cells form more frequent but smaller macropinosomes (Figure 2). We speculate that this indicates that the entirety of the PIP_3_ patch is potentially fusogenic and can internalise vesicles by simply folding onto itself rather than requiring a specific mechanism to orchestrate closure and fission at the rim. How this might be achieved mechanistically is unclear, but is reminiscent of the less organised, more ruffle-like macropinosome formation observed in serum stimulated mammalian epithelial cells, or Ras transformed cancer cell lines (Williamson and Donaldson, 2019).

### GEF, GAP and BAR domain interactions each contribute to RGBARG localisation

Localisation of RGBARG at the interface between the cup interior and the protrusive rim is likely to be critical to effectively control the shape and dynamics of these domains during engulfment. This will position its RhoGEF activity where protrusion is promoted and its RasGAP activity where it can restrain expansion of the interior, leading to the organised cup formation observed in Ax2 cells and absent in *RGBARG* ^*–*^ mutants.

To dissect the mechanisms of RGBARG recruitment we tested the effect of deleting each protein domain in turn. To quantify RGBARG enrichment across the cup, linescans from cup tip to tip were normalised to non-protruding regions of the same cell and averaged across multiple cells (Figure 4A and B). GFP-fused to the cyclic AMP receptor (cAR1-GFP) localises uniformly to the plasma membrane and was used as a control (Figure 4C and D). This method confirmed RGBARG-GFP was enriched 3-fold at the protruding edges of macropinocytic cups and allowed us to quantify how each protein domain contributes to recruitment at the cup (Figure 4 and supplement).

**Figure 4:**
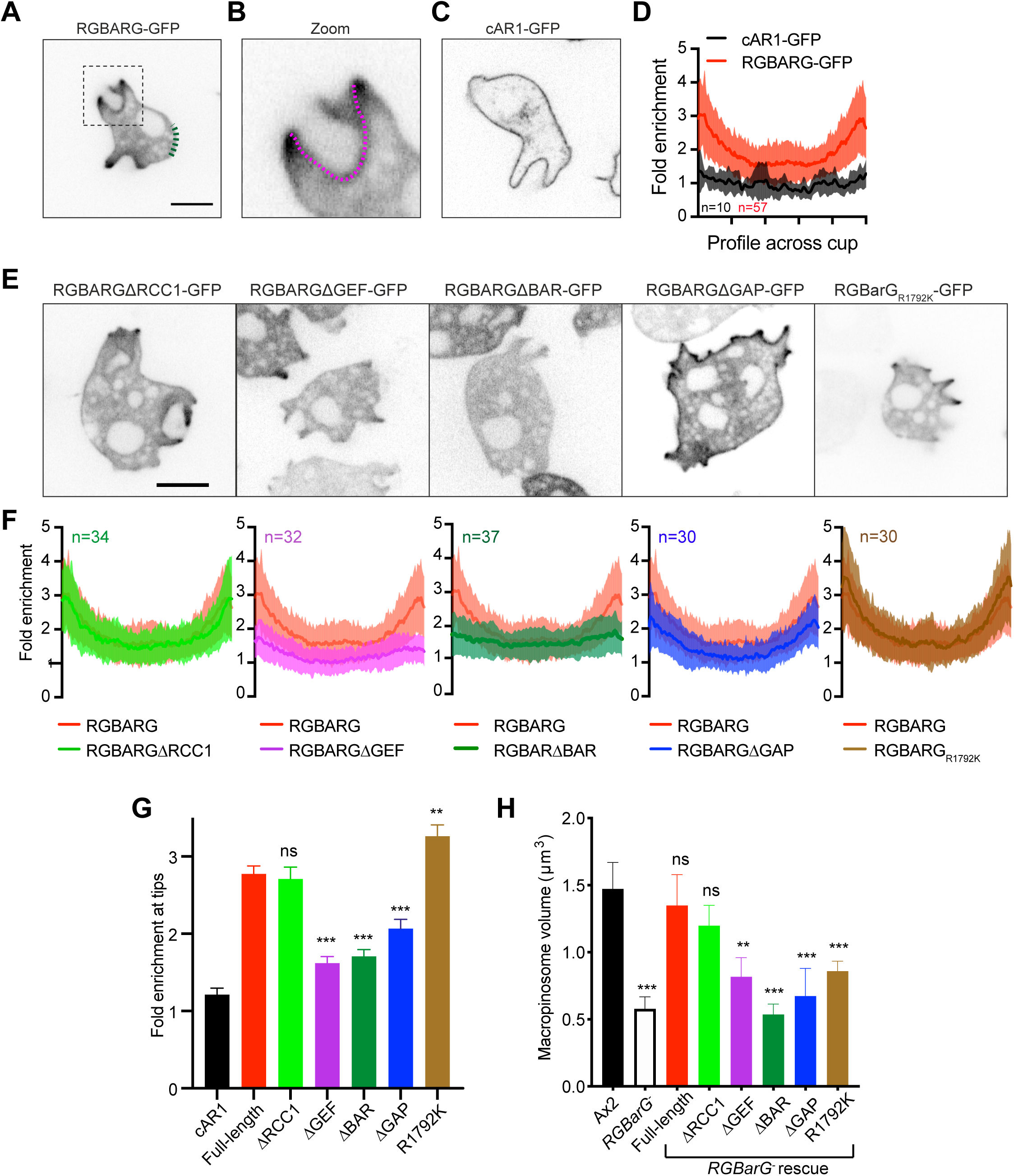
Multiple interaction regulate RGBARG recruitment at the cup. Full-length RGBARG or mutants lacking each domain in turn were expressed as GFP-fusions in RGBARG-cells. (A) Shows an example of full-length RGBARG-GFP. Enrichement was measured relative to the average intensity of a non-protrusive membrane region (green dotted line). The boxed region is enlarged in (B), showing an example of the line measured from the rim along the cup interior. (C) Shows the uniform localization of cAR1-GFP used as a control. (D) Averaged, normalized linescans along the cup from multiple cells, demonstrating a 3-fold enrichment of RGBARG-GFP at the cup rim and uniform cAR1-GFP concentration. (E) Shows representative images of cells expressing RGBARG-GFP with the domains indicated delected, as well as the R1792K point mutation that inactivates RasGAP activity. The averaged intensity of each construct across the cup is shown in (F), with the profile of the full-length protein from (D) in red for comparison. Values plotted are the mean ± standard deviation. (G) The enrichment at the protruding rim of each construct measured by averaging the first 10% of each individual linescan and calculating the mean and SEM across each group. (H) The ability of each construct to rescue large macropinosome formation in RGBARG-cells was determined by measuring the size of nascent FITC dextran-containing macropinosomes by confocal microscopy, as in Figure 2A. >100 macropinosomes over 3 independent experiments were measure. Bars denote mean macropinosome volume ± SEM, ** P<0.01, *** P<0.005 Mann-Whitney T-test.

Removal of the RCC1 domain had no effect on localisation and was able to fully rescue the ability of *RGBARG*^*-*^ cells to form large macropinosomes (Figure 4E-H). In contrast, deletion of either the RhoGEF or BAR domains caused RGBARG to become uniformly cytosolic and did not rescue (Figure 4E-H). In the absence of the RasGAP domain however, RGBARG was still recruited to the plasma membrane but was much more broadly distributed throughout the cup and significantly less enriched at the protruding rim (Figure 4G). RGBARGΔGAP-GFP was also unable to rescue macropinosome formation (Figure 4H). Co-expression of PH_CRAC_-RFP confirmed that RGBARGΔRasGAP-GFP was no longer excluded from PIP_3_, and therefore active Ras domains (Figure 4 supplement 2A). RasGAP interactions therefore restrict RGBARG to the periphery of the cup interior domain.

To confirm the role of the RasGAP interactions in restricting RGBARG localisation, we also made a point mutant in the conserved arginine responsible for stabilising the transition from Ras-GTP to Ras-GDP (Bos et al., 2007). This mutation (R1792K) is predicted to disrupt GAP activity but still allow Ras binding and completely rescued RGBARG exclusion from the cup interior, and slightly but significantly increased enrichment at the cup tip (Figure 4F-G and supplement 2B). However, despite localising to the protruding rim, RGBARG_R1792K_-GFP did not rescue the cup organisation of *RGBARG-*cells, which still produced enlarged PIP_3_ patches and small macropinosomes (Figure 4H and Supplement 2). RGBARG is therefore an active RasGAP and this domain also provides spatial information to position RGBARG to the periphery of the active Ras/PIP_3_ patch where it can prevent its expansion and shape the forming cup.

The data above show that RGBARG integrates spatial cues from its RhoGEF, BAR and RasGAP domains for correct positioning and cup organisation. To identify the relevant binding partners and contribution of each domain, we also expressed them individually fused to GFP. Whilst RhoGEF-GFP expressed too poorly to observe its localisation, both the RCC1 and GAP domains were completely cytosolic (Figure 5A and C). In contrast, the BAR domain alone was sufficient for strong recruitment throughout the plasma membrane (Figure 5B). This was blocked by including either of the adjacent RhoGEF or RasGAP domains (Figure 5D and E), suggesting that BAR-domain binding may also be regulated by intramolecular interactions. In contradiction of our initial hypothesis however, BAR-GFP was not enriched at areas of curvature or protrusion. The BAR domain therefore appears to drive general recruitment to the plasma membrane rather than recognising curvature at cups.

**Figure 5:**
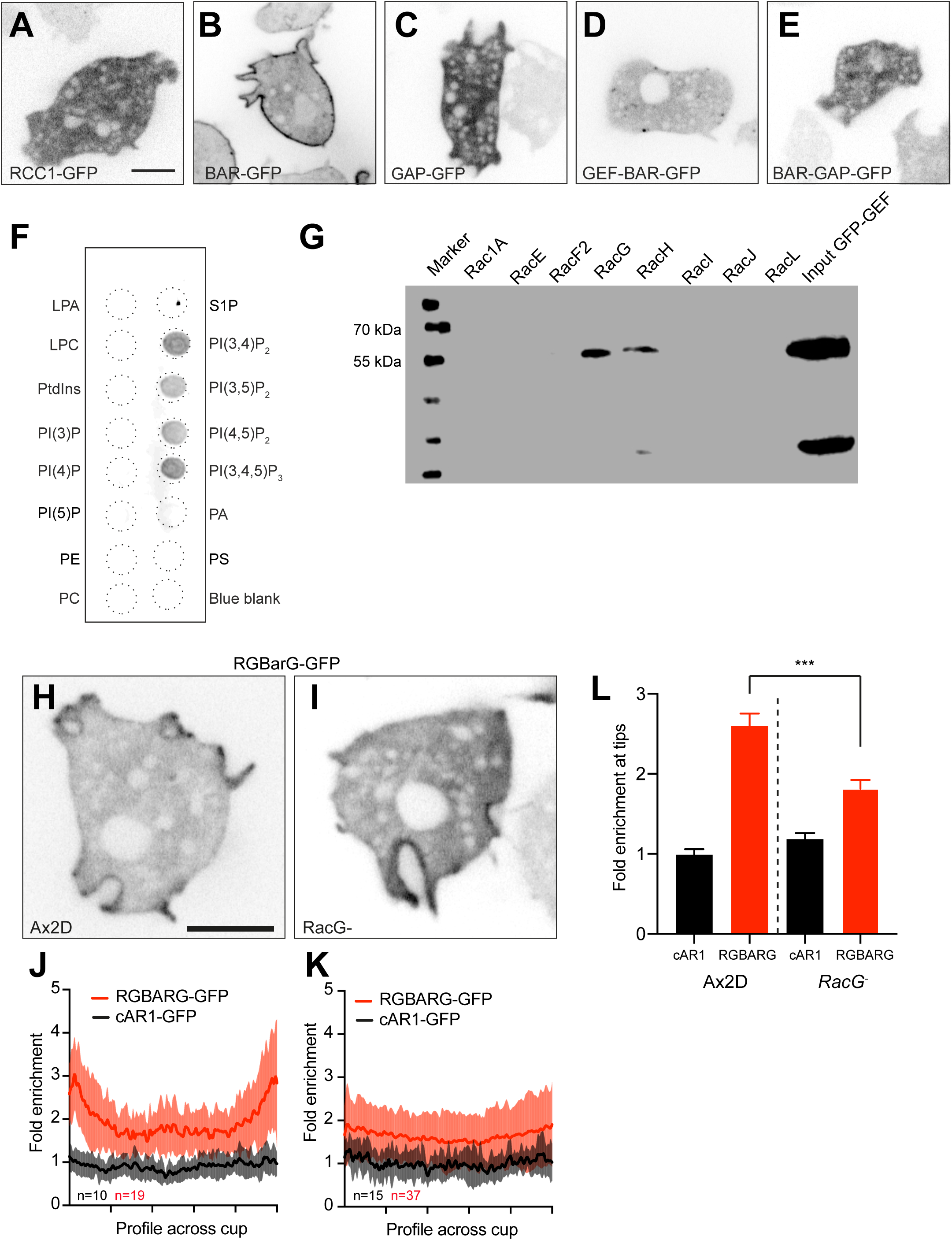
Binding specificity of the BAR and GEF domains. (A-E) The indicates individual, or combination of domain from RGBARG were expressed as GFP fusions in RGBARG-cells. Images shown are single confocal sections. (F) Lipid binding specificity of BAR-GEF by lipid overlay assay using whole cell lysate from BAR-GFP expressing cells. (G) Rac binding specificity of the RhoGEF domain determined by co-immunoprecipitation of GEF-GFP by a library of purified GST-Rac’s bound to beads. (H) and (I) Confocal images of full-length RGBARG-GFP localization in RacG-cells and their parental background strain Ax2D. The average profile (± standard deviation) of RGBARG-GFP along the cup relative in Ax2D and RacG-cells relative to cAR1-GFP is shown in (J) and (K) respectively. (L) Enrichment of RGBARG-GFP and cAR1 at the cup tip in each cell line. Bars indicate mean ± SEM, *** P<0.005 Mann-Whitney T-test. All scale bars indicate 5 μm.

As the BAR domain of RGBARG does not concentrate at specific membrane shapes, we investigated its lipid binding specificity by lipid-protein overlay. Incubation of lysates from cells expressing BAR-GFP with PIP strips indicated binding to all PIPs with two or more phosphates (Figure 5F). This was confirmed by PIP array, indicating a slight selectivity for PI(3,4)P_2_ (Figure 5 Supplement 1A). Given this broad ability to bind all highly phosphorylated phosphoinositides it is likely that this BAR domain generally recognises their high negative charge rather than specific phosphate configurations. This supports a mechanism whereby highly-phosphorylated PIPs recruit RGBARG to the plasma membrane via its BAR domain where additional interactions with the RhoGEF and RasGAP domains further restrict its position and activity to the protruding edges of forming cups.

To identify the targets of the RhoGEF domain, we performed co-immunoprecipitations with a library of recombinant GST-tagged small GTPases. The *Dictyostelium* genome contains an expanded set of Rac small GTPases, but no Rho or CDC42 subfamily members (Vlahou and Rivero, 2006). Of these only RacH and RacG bound the RhoGEF domain of RGBARG with no detectable binding to other Racs, including Rac1 which has previously been implicated in cup formation (Dumontier et al., 2000).

Whilst RacH is involved primarily in endocytic trafficking and localises exclusively to intracellular compartments (Somesh et al., 2006a), RacG localises to the plasma membrane and is enriched at the protruding rim of phagocytic cups (Somesh et al., 2006b). Previous studies have also shown that overexpression of wild-type or constitutively active RacG also promotes phagocytosis, indicating a potential interaction with RGBARG (Somesh et al., 2006b).

Consistent with previous reports, we found loss of RacG had no significant effect on macropinocytosis, with mutants forming normal sized active Ras patches and macropinosomes (Somesh et al., 2006b)(Figure 5 supplement 1B and C). When we measured RGBARG-GFP recruitment, its association with cups was more uniform and was only enriched 1.8±0.6 fold at the rim in *RacG-*cells compared to 2.6±0.7 fold isogenic controls (Figure 5H-L). This indicates that RacG and RGBARG functionally interact *in vivo* and partly contribute to RGBARG localisation. However, the remaining signals or functional redundancy with other Racs is sufficient for partial RGBARG recruitment and apparently normal engulfment in the absence of RacG.

Combined, our data indicate that RGBARG uses a coincidence detection recruitment mechanism to direct cup formation: BAR domain binding to negatively charged phospholipids directs the protein to the plasma membrane whilst additional interactions with RacG and active Ras synergise to constrain RGBARG to the cup rim. This tripartite regulation ensures that RGBARG is accurately positioned to exert its RhoGEF and RasGAP activities at the interface between cup interior and protrusion and organise engulfment.

### RGBARG is a highly active dual specificity Ras/Rap GAP

Forming a complex 3-dimensional shape is likely to require multiple regulators and RGBARG is not the only RasGAP involved in macropinocytosis in *Dictyostelium*. Axenic strains (including the Ax2 parental strain used in this work) also harbour mutations in *NF1* that enhance fluid uptake via enlarged active Ras patches and subsequent formation of larger macropinosomes (Bloomfield et al., 2015). However, whilst mutations in *NF1* were reported in each of 10 independently isolated axenic strains, RGBARG mutations were completely absent ((Bloomfield et al., 2015) and G. Bloomfield, personal communication, May 2019). Our attempts to generate *RGBARG* mutants in non-axenic strains were also unsuccessful. This indicates that disruption of *NF1* but not *RGBARG* is sufficient to support axenic growth, and that they serve different functions.

To better understand the differences between NF1 and RGBARG, we compared the specificity and activities of their RasGAP domains. The *Dictyostelium* genome encodes 14 Ras subfamily members of which RasB, RasG and RasS are the most important for macropinocytosis (Chubb et al., 2000; Hoeller et al., 2013; Junemann et al., 2016; Khosla et al., 2000). Overexpression of RasD can also partially compensate for loss of RasG and S (Khosla et al., 2000). The small GTPase Rap, a close relative of Ras, has also been implicated in macropinosome formation (Inaba et al., 2017). We therefore measured the specific GAP activity from both NF1 and RGBARG against each small GTPase.

Consistent with the inability of RGBARG_R1792K_ to rescue the knockout phenotype, we found that the RasGAP domain of RGBARG was highly active against all the GTPases tested (Figure 6). The RasGAP domain of NF1 was also active against each Ras tested, but with 75% less activity than RGBARG in each case. RGBARG is therefore a more potent RasGAP *in vitro*, but the lack of specificity for particular Ras isoforms for both RGBARG and NF1 indicates their functional differences are likely imparted by additional factors such as their cellular localisation and dynamics.

**Figure 6:**
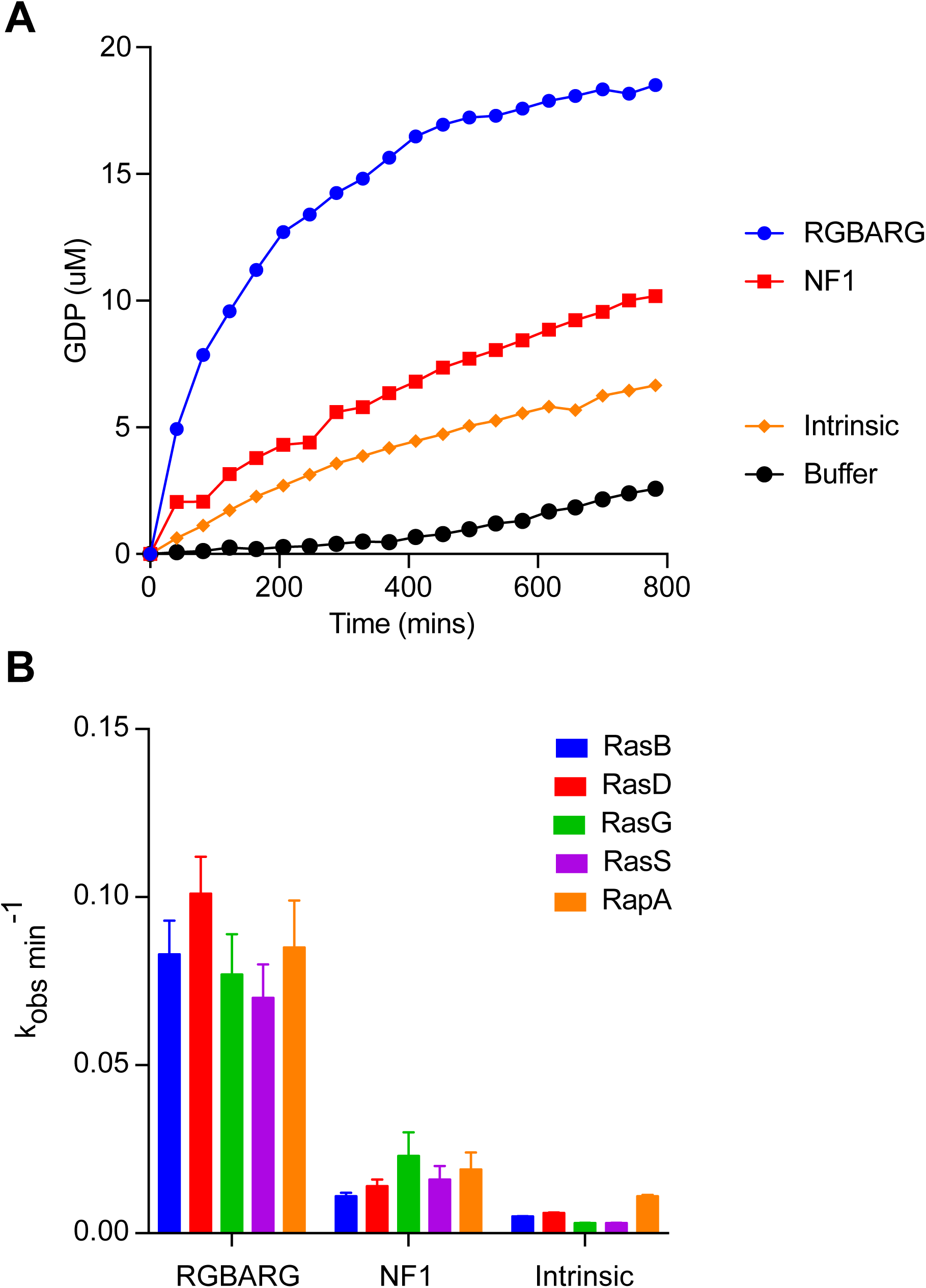
RasGAP activity of RGBARG and NF1. (A) Stimulation of GDP released from GTP-loaded RasG upon addition of recombinant RasGAP domains from either RGBARG or NF1, compared to the intrinsic GAP activity of the GTPase or GTP in buffer. (B) GAP activity of NF1 and RGBARG against a library of Ras superfamily members. Average of 3 independent experiments performed as in (A) in parallel. Bars indicate mean ± standard deviation.

### Loss of RGBARG improves phagocytosis of large objects

The data above demonstrate that RGBARG is important during the spontaneous self-organisation of macropinocytic cups. As RGBARG also localises to phagocytic cups and engulfment of solid particles such as microbes uses much of the same machinery, we also investigated how RGBARG contributes to phagocytosis.

Disruption of NF1 has previously been shown to increase the size of particles that *Dictyostelium* can engulf (Bloomfield et al., 2015). As RGBARG also affects the size of the PIP_3_ domains that define the cup interior we first tested the ability of *RGBARG*^-^ cells to phagocytose different sized beads. Whilst disruption of *RGBARG* had no effect on phagocytosis of 1 μm diameter beads, engulfment of 4.5 μm beads was enhanced 3-fold, with an average of 2.2±0.4 beads engulfed per cell after 1 hour, compared to 1.0±0.4 in control (Figure 7A and B). Enhanced Ras activation therefore appears to be generally beneficial for the engulfment of large spherical targets.

**Figure 7:**
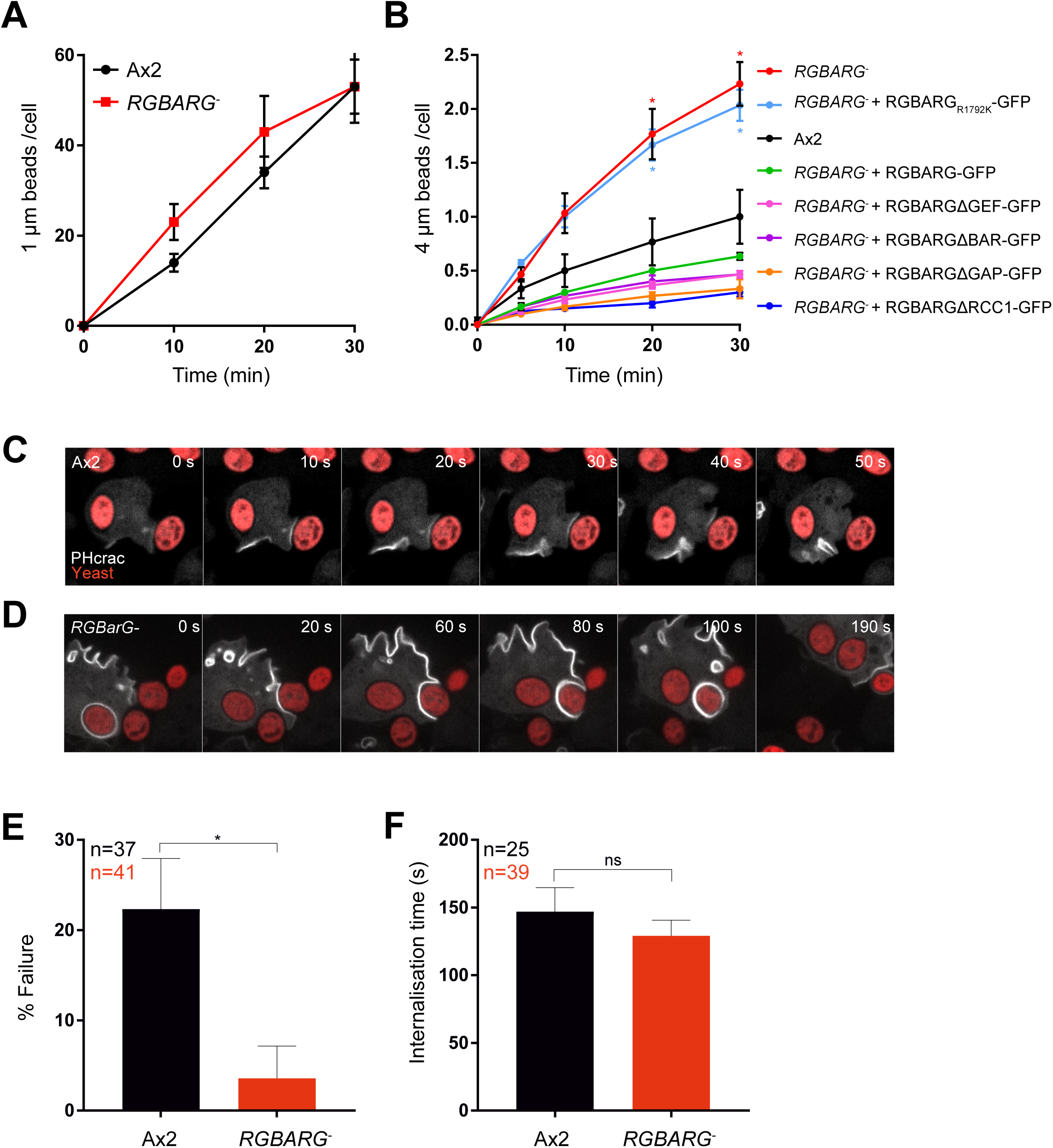
Phagocytic defects in RGBARG-cells. (A) Ax2 and RGBARG-cell have comparable rates of phagocytosis of fluorescent 1 μm beads. (B) Phagocytosis of 4 μm beads by Ax2 cells, RGBARG-cells, or RGBARG-cells expressing full-length or mutant RGBARG-GFP. Phagocytosis measured by flow cytometry in 3 independent experiments. Phagocytosis of TRITC-labelled yeast was directly observed by spinning disc confocal microscopy by either (C) Ax2 or (D) RGBARG-cells expressing PHCRAC-GFP. (C) Shows an example of a failed engulfment. (E) The relative frequency of phagocytosis failure after cup formation (indicated by PHCRAC-GFP recruitment). The time from initial contact to complete engulfment in successful phagocytic events is shown in (F). n indictates total number of phagocytic events over 3 independent experiments. All values plotted are mean ± standard deviation. * P<0.01 Students T-test.

Surprisingly, although expression of RGBARG-GFP from an extrachromosomal vector fully rescued macropinosome formation (Figure 2A-C), this reduced the ability of *RGBARG*^*-*^ cells to engulf 4.5 μm beads to 63% of control levels (Figure 7B). This effect was even more severe upon expression of domain deletion mutants including the ΔBAR, ΔGEF and ΔGAP constructs which do not localise properly and have no deleterious effect on macropinosome formation. This indicates a dominant negative effect, most likely due to sequestration of binding partners. In contrast, expression of RGBARG_R1792K_ had no inhibitory effect on *RGBARG*^-^ cells. This only differs from RGBARG-GFP in its RasGAP activity indicating that mislocalisation or overexpression of this domain is sufficient to inhibit engulfment of large targets.

To better understand how loss of RGBARG affects phagocytosis we performed time-lapse microscopy of cells expressing PH_CRAC_-GFP engulfing TRITC-labeled yeast. Engulfment occurred rapidly in both cell types but failed at a frequency of ∼20% in Ax2 cells with the PIP_3_ patch dissipating and the yeast escaping from the cell (Video 6). Whilst the time for successful engulfment was no significantly altered by loss of *RGBARG* (129±11 minutes in mutants vs 147±18 minutes in Ax2), capture was much more robust with a failure rate of only 4±5 % compared to 22± 7% (Figure 7C-F and Video 7). The main influence on the phagocytic efficiency of large targets in *RGBARG*^*-*^ cells thus appears to be increased cup stability and enlarged Ras signaling patch rather than rate of protrusion around the object.

### Spatial regulation of Ras by RGBARG is important for phagocytosis of elongated targets

To be effective, phagocytic cells must be able to engulf microbes with differing physical properties such as shape, size, stiffness and surface chemistry. As RGBARG is important for the organisation and stability of phagocytic and macropinocytic cups, we also investigated its role during the engulfment of different bacteria.

Phagocytosis was measured by the ability of *Dictyostelium* cells to reduce the turbidity of a bacterial suspension over time. Whilst disruption of *RGBARG* had no effect on the ability to clear a suspension of *Klebsiella aerogenes*, engulfment of *Escherishia coli* was substantially reduced (Figure 8A and B). This was fully rescued by re-expression of RGBARG-GFP. Therefore, although loss of RGBARG has no effect on the engulfment of 1 μm beads and is beneficial for the uptake of large beads and yeast, it causes a specific defect in engulfment of some bacteria.

**Figure 8:**
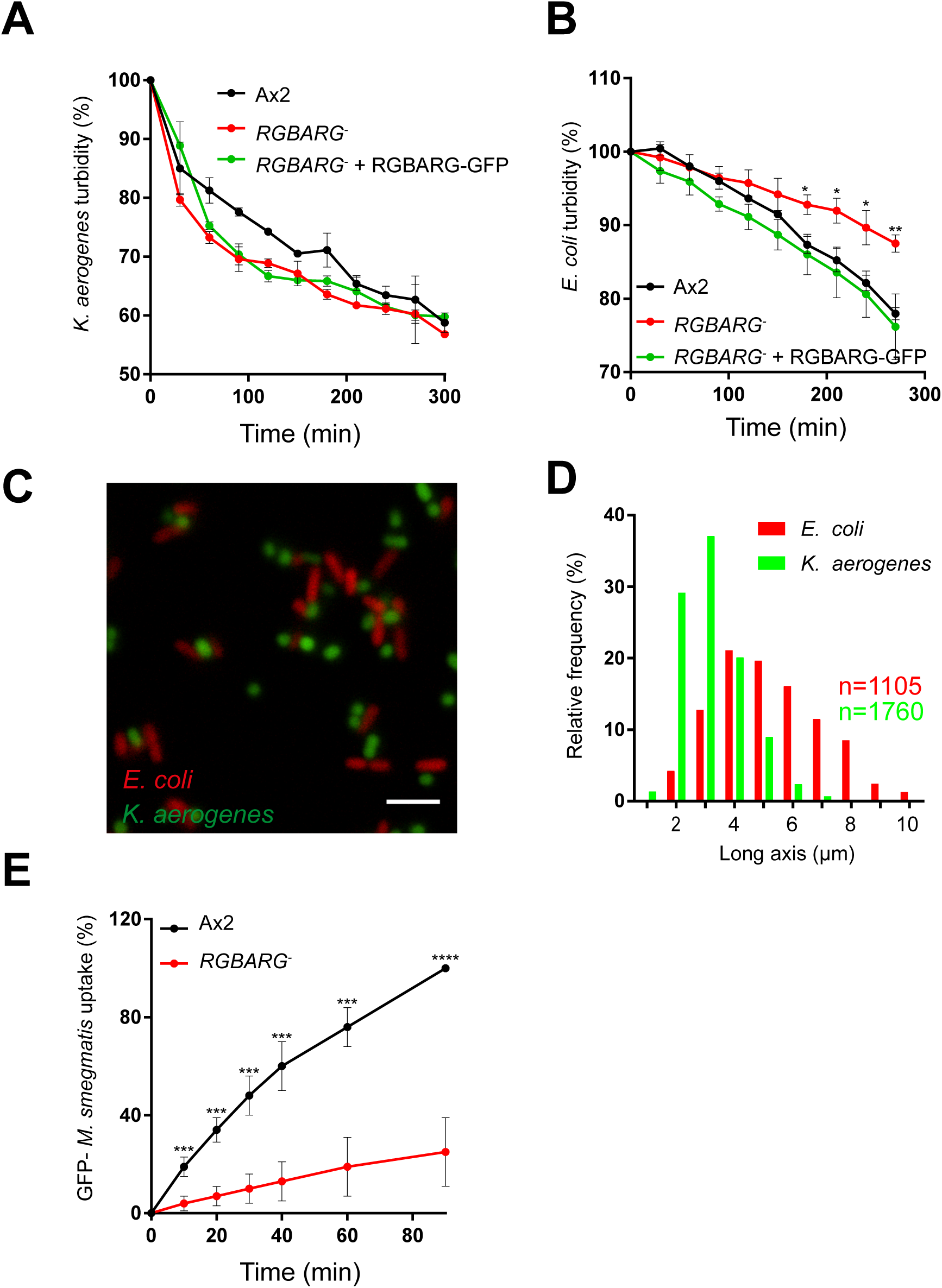
RGBARG-cells have selective defects in phagocytosis of bacteria. Phagocytosis of (A) K. aerogenes or (B) E. coli was measured by monitoring the decreasing turbidity of a bacterial suspension after addition of Dictyostelium. (C) show wide-field fluorescence microscopy images of GFP-expressing K. aerogenes mixed with RFP-expressing E. coli, demonstrating their different shape and size. The length of each bacteria was quantified automatically and is plotted in (D). (E) Phagocytosis of GFP-M. smegmatis, an alternative elongated bacteria, measured by flow cytometry. Values plotted in (A), (B) and (E) are mean ± standard deviation of 3 independent experiments. *P<0.05, **P<0.01, ***P<0.005, Student’s T-test.

**Figure 9:**
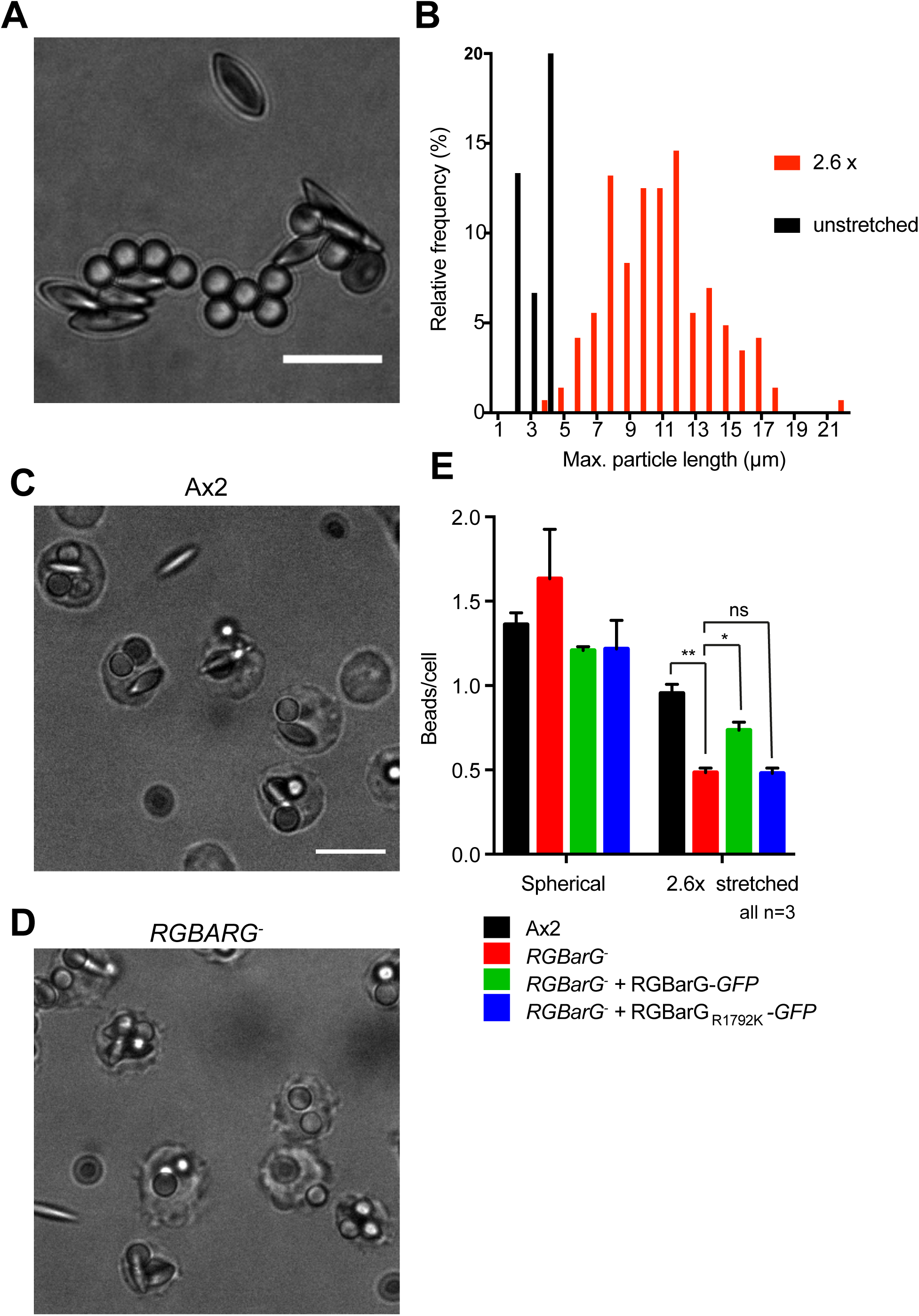
RGBARG-cells have a geometry-specific phagocytic defect. (A) wide-field image of a 50:50 mix of untreated and 2.6-fold stretched polystyrene beads. (B) Quantification of the degree of deformation of stretched beads measured by the long axis. (C) Ax2 and (D) RGBARG-cells were incubated with a 50:50 mix of stretched and unstretched beads for 30 minutes and widefield images acquired. (D) Quantification of the numbed of spherical and ellipsoid beads within each cell showing RGBARG-cells are defective in engulfing the elongated beads. Values plotted are the mean ± standard deviation from 3 independent experiments. * P<0.05, **P<0.01 Mann-Whitney T-Test. All scale bars indicate 10 μm.

The most obvious physical difference between *K. aerogenes* and *E. coli* is their shape (Figure 8C and D). Both have similar short axes but *K. aerogenes* average 3.2 μm in length whilst *E. coli* have an average long axis of 5.4 μm. Previous work investigating phagocytosis of different shaped beads by macrophages concluded that complex elongated shapes are more difficult to engulf (Champion and Mitragotri, 2006). We therefore hypothesised that the selective phagocytosis defects of *RGBARG*^*-*^ cells was due to the target shape. To test this, we measured the ability of *RGBARG*^*-*^ cells to engulf an additional elongated rod-shaped bacteria (GFP-expressing *Mycobacteria smegmatis*, 3-5 μm long) by flow cytometry. The ability of *RGBARG*^*-*^ cells to engulfing these bacteria was again reduced by 75% (Figure 8E), again correlating with an inability to phagocytose elongated targets.

The data above are consistent with a role for RGBARG in enabling the engulfment of elongated bacteria. However, the bacterial strains used will also differ in other aspects such as their surface components, phagocytic receptor activation and stiffness. To directly test the importance of RGBARG in engulfing targets of different shape we therefore stretched 3 μm latex beads to generate oblate ellipsoids of conserved volume and surface chemistry (Ho et al., 1993).

To measure relative phagocytosis in the same experiment, cells were incubated with a 1:1 mix of spherical and stretched beads (2.6x aspect ratio) and the number of engulfed beads of each shape quantified by microscopy. The ability of Ax2 cells to engulf ellipsoid particles was reduced by 30% compared to spheres (P<0.01, T-test). However, whilst *RGBARG*^*-*^ cells engulfed the spheres with similar efficiency, the number of stretched beads taken up was reduced by over 70% (Figure 8C, P<0.01, T-test). These effects were again rescued by re-expression of RGBARG-GFP, but not RGBARG_R1792K_-GFP demonstrating a key role for the RasGAP activity in mediating phagocytosis of elongated particles.

How phagocytic cups organise and adapt their cytoskeleton to engulf targets of differing geometry is very poorly understood. Our data demonstrate that in *Dictyostelium*, a tripartite recruitment mechanism operates to precisely position a RasGAP and RhoGEF domain-containing protein at the interface between the protrusive rim and static interior domains of phagocytic and macropinocytic cups. We propose a model whereby this organises the cup by regulating the balance between protrusion and expansion of the interior in order to both efficiently form macropinosomes and facilitate engulfment of geometrically diverse targets.

## Discussion

In this study we have identified a new component and mechanism used by cells to organise their protrusions into the 3-dimensional cup shapes required to engulf extracellular fluid or particles. Consistent with previous studies, our data support a model whereby cup formation is guided by the formation of a protrusive rim encircling a static interior domain (Veltman et al., 2016). We show that in *Dictyostelium*, RGBARG provides a direct link between the Ras and Rac activities that underlie these different functional domains, providing a novel mechanism to co-ordinate cup organisation in space and time.

RGBARG is not the only RasGAP in *Dictyostelium* involved in macropinosome formation. However, it is the only RasGAP in the *Dictyostelium* genome to also possesses a RhoGEF domain. RGBARG is therefore unique in its ability to integrate the activities of both GTPase families. There are no human proteins with an identical domain structure to RGBARG, and whilst most classical RasGAPs are found in multidomain proteins, none also contain a classical RhoGEF domain (Bos et al., 2007). A screen for RhoGAPs involved in phagosome formation in macrophages identified 3 proteins (ARHGAP12, ARHGAP25 and SH3BP1), but although they all contain PIP_3_ binding (PH) or BAR domains, none contain domains that link to other GTPase families (Schlam et al., 2015). The oncogene TIAM1 contains both RhoGEF and Ras-binding domains however, and BAR domains are found in conjunction with GAP or GEFs in several other proteins. Therefore, whilst mammalian cells also need to coordinate Ras and Rac activity during cup formation, this is likely achieved via multiple proteins, potentially in a complex.

Multiple GAP’s and GEF’s collaborate to shape protrusions into cups. This is apparent in the different roles played by RGBARG and NF1. Both are important negative regulators of Ras, but whilst RGBARG is specifically enriched at the cup rim, NF1 appears to be present thoughout the cup (Bloomfield et al., 2015). RGBARG and NF1 therefore play different functional roles; whilst disruption of NF1 leads to an increase in the volume of fluid taken up and can facilitate axenic growth (Bloomfield et al., 2015), RGBARG appears more important for cup structure and shape. We therefore speculate a model whereby NF1 acts to generally suppress Ras activity and regulate the spontaneous excitability of active Ras patches, whilst RGBARG operates at their periphery to restrict their expansion and stimulate protrusion via Rac. This is illustrated in Figure 10.

**Figure 10:**
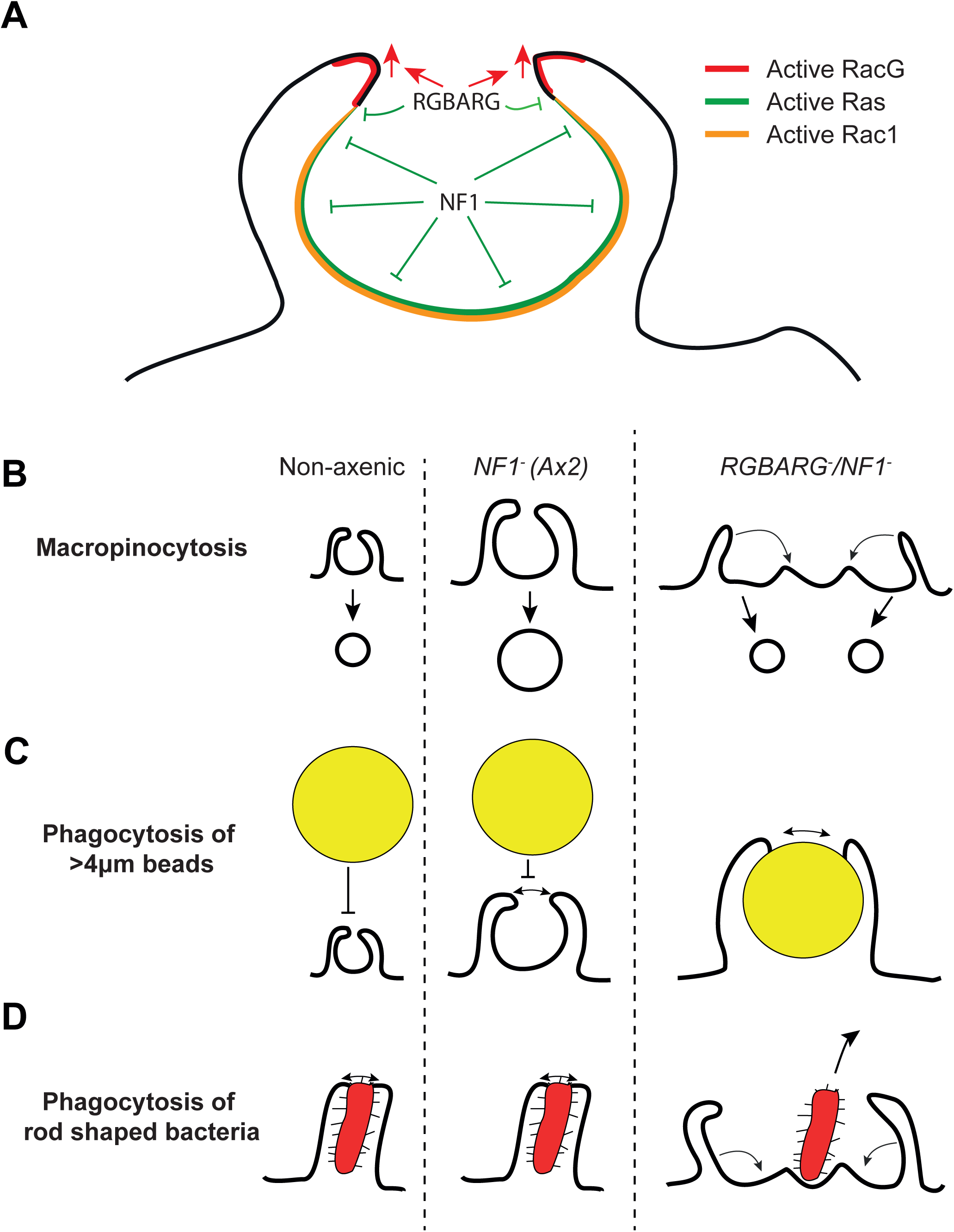
Model of RGBARG function. (A) RGBARG functions specifically at the protrusive rim of the cup and simultaneously activates protrusion by activation of RasG, whilst locally inhibiting Ras activity to restrict expansion of the cup interior. In contrast, NF1 is present throughout the cup interior, playing a more general role in suppressing Ras activity. Below are cartoons depicting the effects of disrupting NF1 and RGBARG on (A) macropinosome formation, (B) phagocytosis of large objects and (C) phagocytosis of elongated bacteria. Whilst loss of NF1 appears to simply make cups larger, subsequent disruption of RGBARG causes wide, flat ruffles that form multiple small macropinosomes. The enlarged size of these ruffles assists engulfment of large particles, but is detrimental to the uptake of more complex shapes such as elongated bacteria or beads.

This model is doubtless overly simplistic, and other RasGAP’s also contribute to shaping active Ras dynamics. For example, the IQGAP-related protein IqgC was also recently shown to have RasGAP activity and localise throughout the interior of macropinocytic and phagocytic cups in *Dictyostelium* (Marinovic et al., 2019). In contrast to our findings with NF1 and RGBARG however, IqgC is reported to have specific RasGAP activity against RasG. As the different Ras isoforms are non-redundant (Khosla et al., 2000), IqgC adds a further layer of regulatory complexity shaping the dynamics of engulfment.

Whilst the regulation of Ras signaling and the static interior domain is becoming clearer, how protrusion is regulated during engulfment is less well understood. In mammalian cells, several studies indicate that actin dynamics and protrusion at the cup is regulated by the combined activities of Rac1 and CDC42 (Cox et al., 1997; Massol et al., 1998; Schlam et al., 2015). Rac1 and CDC42 are differentially activated with active Rac1 throughout the cup and CDC42 activation earlier and more restricted to the rim (Hoppe and Swanson, 2004). Whilst *Dictyostelium* does not possess a clear CDC42 orthologue, we find that RGBARG specifically interacts with the atypical Rac isoforms RacG and RacH. Currently, no direct effectors of either protein are known, and RacG does not interact with the Rac-binding domain of PAK commonly used as a probe for active Rac1 (Somesh et al., 2006b). RacG therefore has at least partly distinct effectors to Rac1. Nonetheless, in cell-free assays, RacG is able to induce actin polymerisation via the ARP2/3 complex, although whether this is dependent on SCAR/WAVE or other WASP family members is not known (Somesh et al., 2006b). RacG therefore appears to be at least partly responsible for defining the protrusive rim, possibly through some coincidence-detection mechanism with active Rac1.

Whilst RacG has no clear direct orthologue in mammalian cells, it is most similar to Cdc42 in protein sequence. Whilst constitutively active Rac1 induces the formation of lamellipodial-type protrusions (Dumontier et al., 2000), constitutively active RacG and Cdc42 both induce filopodia (Nobes and Hall, 1995; Somesh et al., 2006b). We therefore speculate that RacG and Cdc42 are functional orthologues, and the mechanism by which these small GTPases integrate with active Ras to restrict the localisation of RhoGEF and RasGAP proteins such as RGBARG is a general device used to define the protrusive cup rim.

The involvement of both RacG/Cdc42 and Rac1 family GTPases indicates a complex relationship between filopodial and lamellipodial type protrusions during cup formation. Whilst most studies in *Dictyostelium*, RAW macrophages, dendritic cells and cancer cell lines describe macropinosome formation from smooth, sheet-like projections or cups (Swanson, 2008; Veltman et al., 2016; West et al., 2000; Williamson and Donaldson, 2019), it was recently shown that RAW macrophages can also form macropinosomes by a more filopodial “tent-pole”-type mechanism where protrusion is driven by actin-rich spikes (Condon et al., 2018). Whether this is a general mechanism, or represents a shift in the balance of filopodial vs lamellipodial regulatory proteins in these cells is unclear. However it is probable that filopodial and lamellal cup formation are non-exclusive extremes of a continuum-much as they are during cell migration.

The multi-layered regulation of small GTPases is particularly important when cells are challenged to engulf particles or microbes of different shapes. This is critical for amoebae to feed on diverse bacteria or immune cells to be able to capture and kill a wide range of pathogens, but how cells adapt to different target geometries is very poorly understood (Champion and Mitragotri, 2006; Champion and Mitragotri, 2009). To our knowledge, *RGBARG*^-^ cells are the first mutants described with a geometry-specific defect in phagocytosis, underlining the importance of co-ordinating and balancing Ras and Rac activities. This again differs from the role of NF1, as the *NF1-*deficient Ax2 strain used in this study is able to efficiently engulf and grow on a wide range of bacteria including elongated strains such as *E. coli* (Figure 10) (Buckley et al., 2019). It is still not known how other regulatory elements or cytoskeletal components adapt to differing shapes, but it seems likely that large-scale rearrangements are necessary to accommodate different targets.

In summary, we describe a mechanism to co-ordinate the activity of Rac and Ras family GTPases during engulfment in *Dictyostelium*. The proteins that mediate this co-ordination in mammalian cells remain unknown. However, we propose a general model by which spatial signals and effectors from multiple small GTPases integrate to shape the protrusions that form macropinocytic and phagocytic cups, enabling cells to engulf diverse targets.

## Methods

### Dictyostelium culture and molecular biology

Unless otherwise stated, *Dictyostelium* strains were derived from the MRC-Ax2 axenic strain provided by the Kay laboratory and were routinely cultured in filter sterilised HL-5 medium (Formedium) at 22°C. RacG mutants and corresponding parental strain (from the Devreotes group, Johns Hopkins, Ax2D) were kind gifts from Francisco Rivero (University of Hull)(Somesh et al., 2006b). Growth rates were measured by seeding cells at 0.5 × 10^5^/ml in HL-5 and counting cell number twice daily for three days. Growth rate was then calculated by fitting an exponential growth curve using Graphpad Prism software. Cells were transformed by electroporation: 6 × 10^6^ cells were resuspended in 0.4 mls of ice cold E-buffer (10 mM KH_2_PO_4_ pH 6.1, 50 mM sucrose) and transferred to 2 mm electroporation cuvette containing DNA (0.5 µg for extrachromosomal plasmids, 15 µg for knockout vectors). Cells were then electroporated at 1.2 kV and 3 µF capacitance with a 5 Ω resistor in series using a Bio-Rad Gene Pulser II. After 24 hours transformants were selected in either 20 µg/ml hygromycin (Invitrogen), 10 µg/ml G418 (Sigma) or 10 µg/ml blasticidin (Melford).

BAR domain contain proteins were identified by multiple BLAST searches using Dictybase (www.dictybase.org) (Fey et al., 2013). Coding sequences were then amplified by PCR from vegetative Ax2 cDNA adding compatible restriction sites for subcloning into the BglII/SpeI sites of the N- and C-terminal GFP-fusion *Dictyostelium* extrachromosomal expression vectors pDM1043 and pDM1045 (non-axenically selectable versions of the pDM modular expression system (Veltman et al., 2009)). Truncation and point mutants of RGBARG were also generated by PCR and expressed using pDM1045. The *RGBARG* (DDB_G0269934) knockout construct was generated by PCR fusion of ∼1Kb 5’ and 3’ recombination arms with the floxed blasticidin selection cassette from pDM1079, as described in detail in (Paschke et al., 2018). After transformation, independent clones were obtain by dilute plating in 96 well plates. Disruption of the RGBARG locus was screened by PCR from genomic DNA isolated from 1 × 10^6^ cells lysed in 100 μl 10mM Tris-HCl pH8.0, 50 mM KCl, 2.5mM MgCl1, 0.45% NP40, 0.45% Tween 20 and 0.4 mg/ml Proteinase K (NEB). After 5 minutes incubation at room temperature, the proteinase K was denatured at 95°C for 10 minutes prior to PCR analysis. The Ras binding domain (RBD) of PAK1-GFP construct used as an active Ras reporter was a gift from Gareth Bloomfield.

### Macropinocytosis assays

To measure bulk fluid uptake 2mg/ml FITC-dextran (70kDa; Sigma) was added to cells at 5 × 10^6^/ml in shaking culture. At each timepoint, 500μl of cells were removed and added to 1ml ice-cold KK2 (16.5mM KH_2_PO_4_, 3.8mM K_2_HPO_4_, pH6.1). Cells were then pelleted at 7,000 x *g* for 30 seconds, washed once in KK2 and frozen. Pellet were then lysed in 200μl 50mM Na_2_PO_4_ pH9.3, 0.2% Triton X100) and fluorescence measured on a plate reader at 485 excitation/520nm emission. Fluorescence was then normalised to total protein in an additional sample and calculated as a percentage of wild-type cells at 90 minutes.

Macropinosome volume was measured by incubating cells for 5 minutes in 0.1μg/ml FITC-dextran and obtaining Z-stacks on a spinning disc confocal microscope. FITC is pH-sensitive and the sensitivity was set so only new non-acidified macropinosomes were visible. For analysis, individual cells were cropped out, randomised, and volume calculated from manually measuring the maximum diameter of each macropinosome in each cell, assuming they were spherical.

### Phagocytosis assays

Phagocytosis of fluorescent beads was measured by flow cytometry as previously described in detail (Sattler et al., 2013). Briefly, 1 or 4.5 μm diameter YG-carboxylated polystyrene beads (Polysciences Inc) were shaken with 2 × 10^6^ *Dictyostelium* /ml at ratios of 200:1 and 10:1 respectively. 500 μl samples were removed at each timepoint and added to 3 ml ice-cold Sorenson sorbitol buffer (SSB; 15 mM KH_2_PO_4_, 2 mM Na_2_HPO_4_, 120 mM Sorbitol) containing 5mM sodium azide. Samples were then centrifuged at 100 x *g* for 10 minutes, pellets resuspended in SSB and analysed on an Attune NxT flow cytometer (Life Technologies). Analysis was performed using FloJo software as described (Sattler et al., 2013).

To measure uptake of GFP-expressing *M. smegmatis* by flow cytometry, the bacteria were grown to an OD_600_ of 1, pelleted by centrifugation at 10,625 x g for 4 minutes and resuspended in 1ml HL5 medium. Bacteria clumps were then disrupted by passing through a 26-guage needle several times, before adding a 1/10th volume of bacteria to a *Dictyostelium* culture and processing as above.

To measure phagocytosis of bacteria by decreasing turbidity, an overnight bacterial culture in LB was diluted 1:25 and grown at 37°C until an OD_600_ of 0.7 before pelleting and resuspension in SSB at an OD_600_ of 0.8. This was then added to an equal volume of *Dictyostelium* at 2 × 10^7^ cells/ml in SSB at room temperature and shaken in flasks. The OD_600_ was then measured over time.

Phagocytosis and TRITC labeling of heat killed *S. cerevisiae* was performed essentially as previously described (Rivero and Maniak, 2006). *Dictyostelium* at 1 × 10^6^ cells/ml in HL5 were seeded in glass-bottomed microscopy dishes (Mat-tek) and left for 1 hour prior to addition of a 5-fold excess of yeast. After 30 minutes, the fluorescence of extracellular yeast was quenched by addition of 0.2 mg/ml trypan blue and images of multiple fields of view taken on a wide-field microscope scoring >100 cells per condition.

### Microscopy and image analysis

Live cell imaging was performed in glass-bottomed microscopy dishes (Mat-Tek) with cells seeded the preceding day in filtered HL-5 medium, unless otherwise stated. Spinning disc images were captured using a Perkin-Elmer Ultraview VoX spinning disk microscope with a UplanSApo 60x oil immersion objective (NA 1.4) and Hammamatsu C9100-50-EM0CCD camera. Laser scanning confocal images were obtained using a Zeiss LSM880 Airyscan confocal equipped with a Fastscan detector, and a 63x 1.4 NA objective. Images were acquired in fastscan mode and deconvolved by airyprocessessing using Zen black software (Zeiss).

Image analysis was performed using ImageJ (https://imageJ.nih.gov) with average plots of protein enrichment across cups generated using a custom script in Igor Pro (Wavemetrics) (see Supplement to methods). For this, confocal images were captured and linescans of GFP-fluorescence intensity measured from the protrusive tip to tip. The average signal from a 1-2 μm non-protruding region of the cell was also measured as was the local background outside each cell. Local background was subtracted and signal across the cup divided by the non-protruding membrane signal to give fold-enrichment. To compare enrichment across multiple cups, normalised linescans were extrapolated over 1000 points and averaged. Enrichment at the cup tip was measured by the average of the first 100 points of the profile for each cup.

The bacterial long axis was measured automatically from widefield images of either GFP or RFP expressing bacteria in ImageJ. Individual bacteria were identified by thresholding and long axis measured using the Feret’s diameter function.

### Western blotting and Lipid overlay assays

Western blotting was performed by standard techniques, separating proteins by SDS-PAGE and probing using a custom rabbit polyclonal antibody to GFP (gift from Andrew Peden). Endogenously biotinylated proteins were used as a loading control, using Alexa680-labelled streptavidin (Life Technologies)(Davidson et al., 2013). Blots were imaged LiCor odyssey SA fluorescence gel imager.

For lipid overlay assays, 1 × 10^7^ Ax2 cells expressing the BAR domain fused to GFP (pCB114) were washed once in SSB, lysed in 600 μl RIPA buffer (50mM TrisHCl pH7.5, 150mM NaCl, 0.1% SDS, 2mM EDTA, 0.5% sodium deoxycholate, 1 x HALT protease inhibitors (Thermo Fisher), 0.5% Triton X100) and left on ice for 45 minutes. Insoluble material was then removed by centrifugation at 15,871 x *g* for 20 minutes at 4°C. PIP strips or Arrays (Echelon Biosciences) were blocked in 3% fatty acid-free bovine serum albumin (BSA) in TBS-T (20mM Tris base, 150mM NaCl, 0.05% Tween20, pH7.2). Samples were then diluted in TBS-T and incubated with the strips for 1 hour at 22°C, before washing and processing as for Western blotting.

### GAP and GEF biochemistry

Interactions with recombinant GST-Rac isoforms were performed as described previously (Plak et al., 2013). *Dictyostelium* cells expressing GST-Rac bait proteins and GFP-fused to the GEF domain of RGBARG were expressed and lysed in 2mls of buffer (10mM Na2HPO4 pH7.2, 1% Triton X100, 10% glycerol, 150mM NaCl, 10mM MgCl_2_, 1mM EDTA, 1mM Na_3_VO_4_, 5mM NaF) including protease inhibitor cocktail (Roche). Lysates were mixed with glutathione sepharose beads (GE Healthcare) and incubated overnight at 4°C. Unbound proteins were washed away with PBS, and bound proteins detected by Western blot using an anti-GFP antibody (SC9996).

For GAP activity measurements, His-NF1 GAP domain (AA 2530-3158) and MBP-His-RGBAR GAP domain (AA 1717-2045) were produced and isolated from *E. coli* Rosetta cells. His-NF1 GAP was purified using a HisTrap excel-affinity column (GE Healthcare) and eluted in buffer containing 50mM Tris, 50mM NaCl, 5% Glycerol, 3mM β-mercaptoethanol and 200mM imidazole, pH7.5. MBP-His-RGBAR GAP, was purified by Maltose Binding Protein Trap (MBPTrap)-affinity column (GE Healthcare) and eluted in 20mM Tris, 200mM NaCl, 5% Glycerol 1mM β-Mercaptoethanol and 10mM Maltose, pH7.5. Proteins were further purified by size exclusion chromatography (Sephacryl 16/60, GE Healthcare) and stored in 50mM Tris, 50mM NaCl, 5mM DTT, and 5mM MgCl2, pH7,5.1.

1µM of the indicated Ras proteins with and without equal amount of indicated GAP domain was incubated with 50 µM of GTP at 20°C in 50 mM Tris pH 7.5, 50 mM NaCl and 5mM MgCl_2_. At each timepoint the GDP content of the samples was analysed by a HPLC (Thermo Ultimate 3000): a reversed phase C18 column was employed to detect GDP and GTP content (in %) as previously described (Eberth and Ahmadian, 2009). Linear rates of GDP production (first 4-8 timepoints) were calculated using GraFit 5.0 (Erithacus software).

### Ellipsoid beads generation and phagocytosis

3µm unmodified non-fluorescent polystyrene beads (Polysciences Inc.) were embedded in polyvinyl alcohol (PVA) film (Sigma Aldrich) and stretched as previously described (Ho et al., 1993). Briefly, 2.8 mls beads of bead solution were added to 20mls 25% w/w dissolved PVA solution and poured into a 10.5 × 10.5 cm plastic mold to create a film. These were cut into 3 × 2cm strips, marked with a grid to follow deformation and placed in a custom stretching device as described in detail in (Ho et al., 1993). Films were the placed in a 145 °C oil bath to soften beads and film and slowly pulled to the desired length. After cooling below the glass transition Tg temperature, the beads were extracted from the central region where the grid was deformed evenly. This part was cut into small pieces and rotated in 10 mls of a 3:7 mix of isopropanol:water overnight at 20°C to dissolve. Beads were aliquoted and twice heated at 75°C for 10 minutes and washed in isopropanol: water. Beads were then washed twice in isopropanol:water at 22 °C, before two washes in water. The amount of stretch was measured by imaging on an inverted microscope and manually measuring their length in ImageJ.

To measure phagocytosis, equal numbers of stretched and unstretched beads were mixed, sonicated and incubated at a 10-fold excess to cells at 1 × 10^6^ /ml, shaking in HL5. After 30 minutes, 500 µl samples were added to 3 ml SSB with 5mM sodium azide to detach unengulfed beads. Cells were washed in ice-cold SSB, transferred to a microscopy dish and allowed to adhere for 10 minutes before imaging and the number of each shape bead internalised quantified manually.

## Acknowledgements

The authors would like to thank Francisco Rivero for providing the RacG mutant cell line and plasmids, Andrew Peden for the GFP antibody, Gareth Bloomfied and Rob Kay for NF1 and GFP-RBD constructs and Iwan Evans for the RFP-*E. coli* strain. JSK is supported by Royal Society University Research Fellowship UF140624. Microscopy studies were supported by UK Medical Research Council grant (G0700091) and Wellcome Trust grant (GR077544AIA). DT was supported by MRC core funding to Rob Kay MC_U105115237.

## Supplementary videos

**Video 1**: RGBARG-GFP is enriched at the tips of macropinocytic cups. Movie of *RGBARG-*cells expressing RGBarG-GFP in axenic culture.

**Video 2**: RGBARG-GFP is enriched at the tips of phagocytic cups. *RGBARG-*cells expressing RGBarG-GFP engulfing a TRITC-labelled budded yeast.

**Video 3**: RGBARG-GFP localises to the periphery of PIP3 patches during macropinocytosis. 3D movie of cells macropinosome formation in *RGBARG-*cells expressing RGBARG-GFP and PH_CRAC_-RFP.

**Video 4:** PIP_3_ dynamics and cup formation in Ax2 cells. Maximum intensity projection of Ax2 cells expressing PH_CRAC_-GFP, showing removal of PIP3 signalling after macropinosome formation is complete.

**Video 5:** *RGBARG-*cells have larger and more persistent PIP3 patches. Maximum intensity projection of cells expressing PH_CRAC_-GFP.

**Video 6:** Phagocytosis of TRITC-labelled yeast by Ax2 cells expressing PH_CRAC_-GFP.

**Video 7:** Phagocytosis of TRITC-labelled yeast by *RGBARG-*cells expressing PH_CRAC_-GFP.

**Figure 2 supplement:**
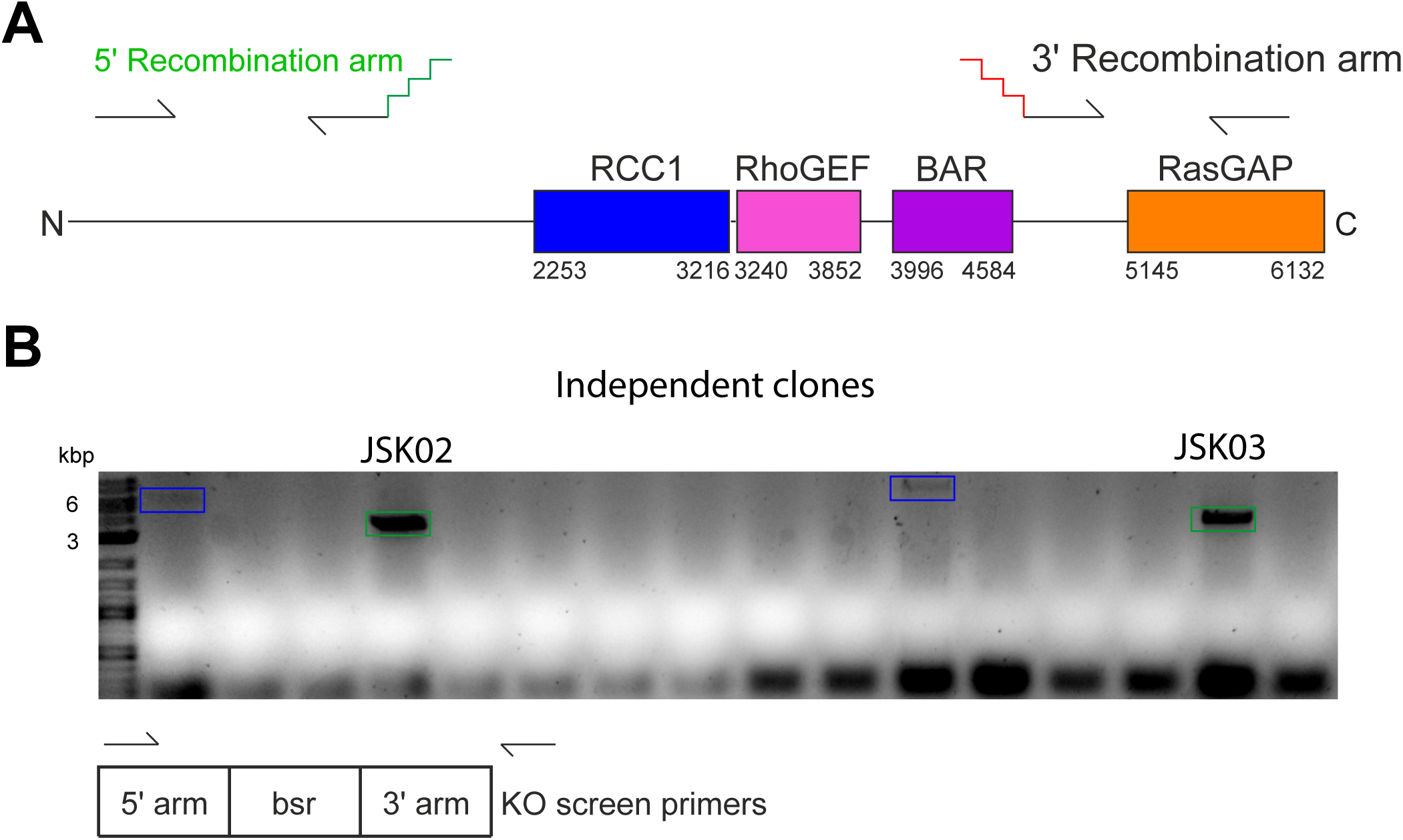
Disruption of DDB_G0269934. (A) Schematic of the genomic locus indicating the position of the regions encoding each domain, and the 5’ and 3’ recombination arms amplified by PCR. These were attached either side of a blasticidin selection cassette by fusion PCR and used to transform Ax2 cells and delete ∼3 kbp of the gene. (D) PCR screen of transformants, using one primer within the 5’recombination arm and another after the 3’ arm. Clones with DDB_G0269934 disrupted will give a product of 3.1 kbp (green box), the wild-type locus is 6.1 kbp (blue boxes).

**Figure 3 supplement:**
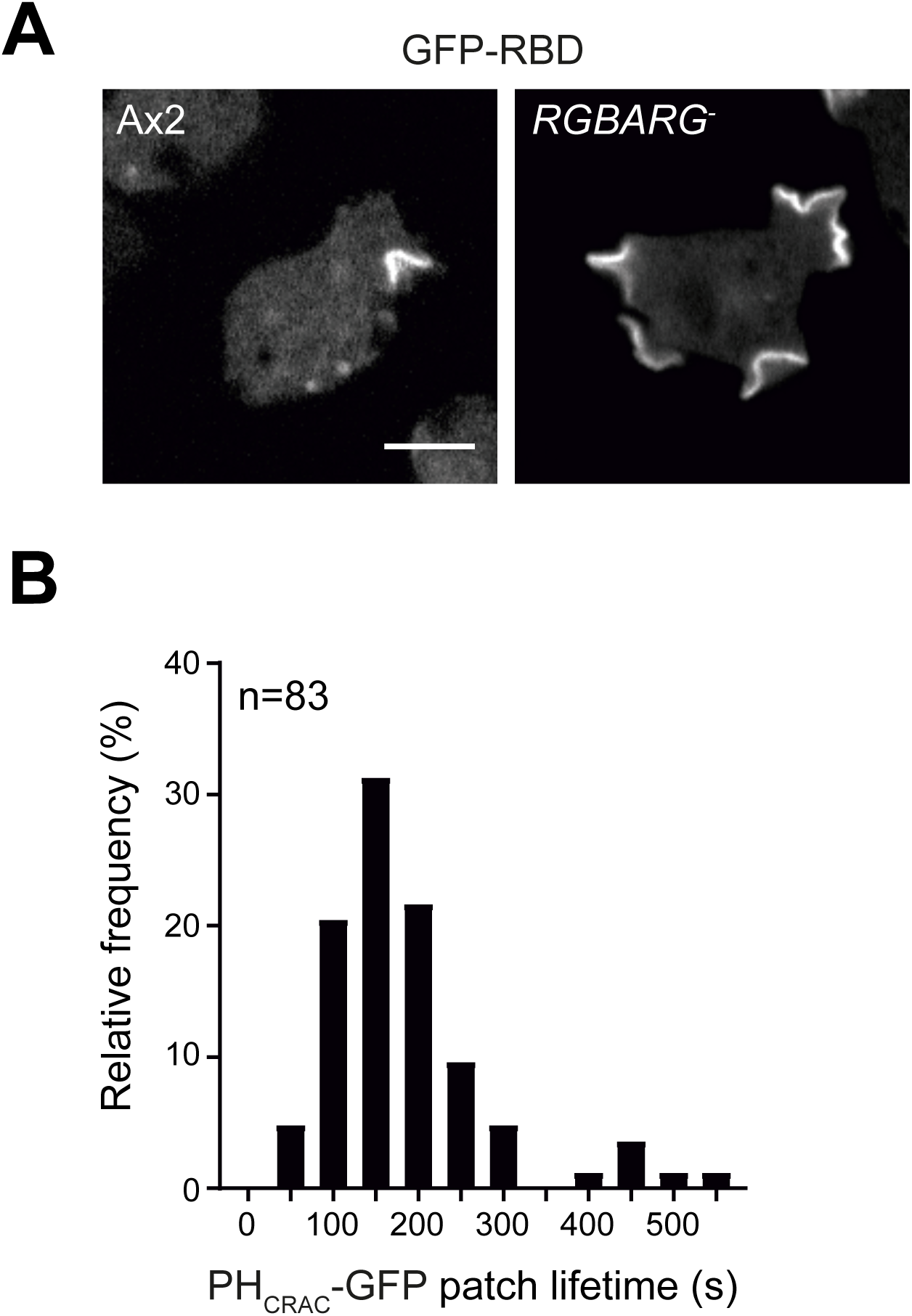
(A) Distribution of the active Ras probe RBD-GFP in wild-type and RGBARG-cells. Images are single confocal planes, bar indicates 5 μm. (B) Histogram of PHCRAC-GFP patch lifetime in Ax2 cells from maximum projection movies. Lifetime was measured from the first frame an independent patch was visible to when it was completely removed from both the surface and any internalized vesicle.

**Figure 4 Supplement 1:**
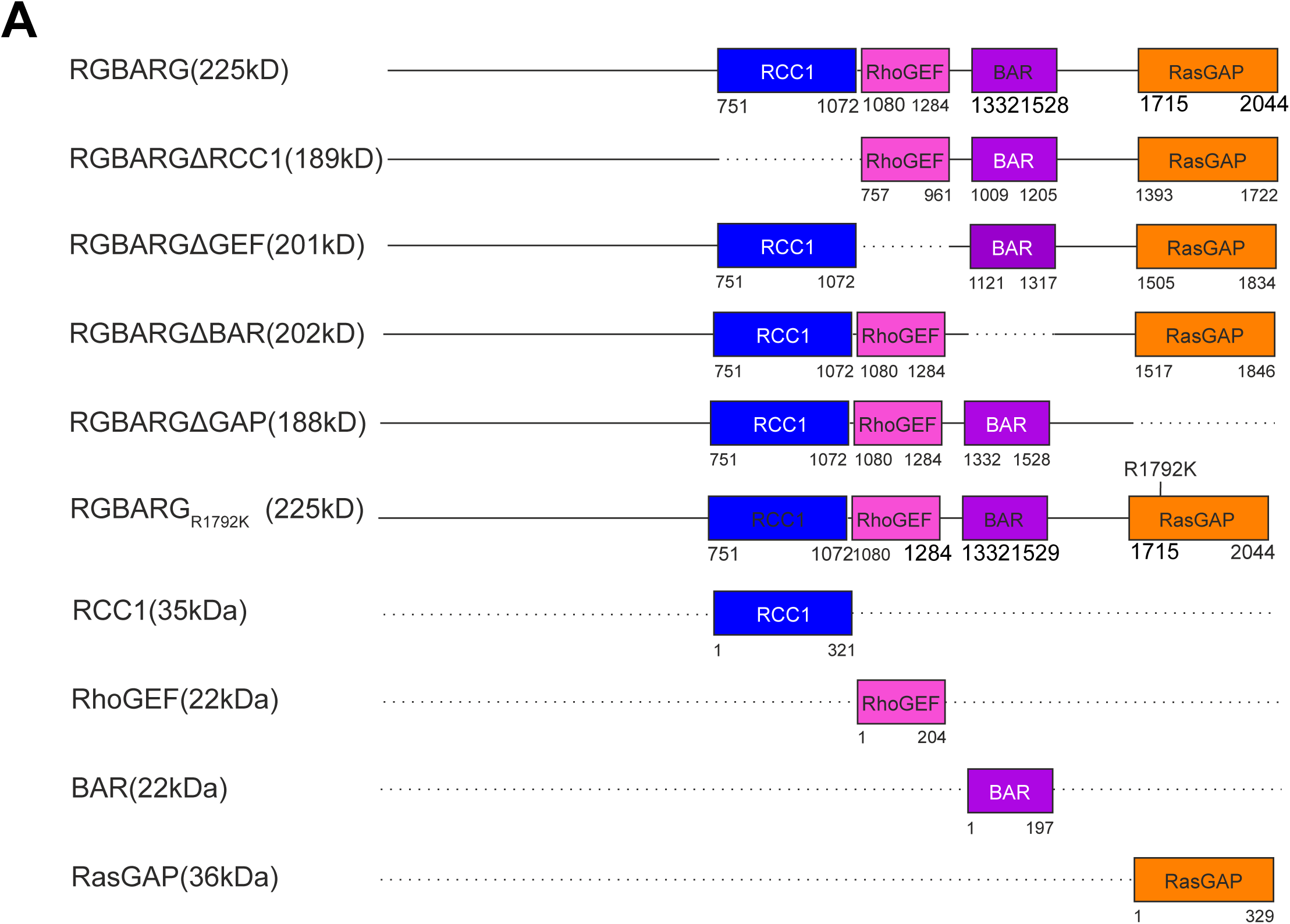
Schematic of the RGBARG truncation and point mutants used in this study.

**Figure 4 Supplement 2:**
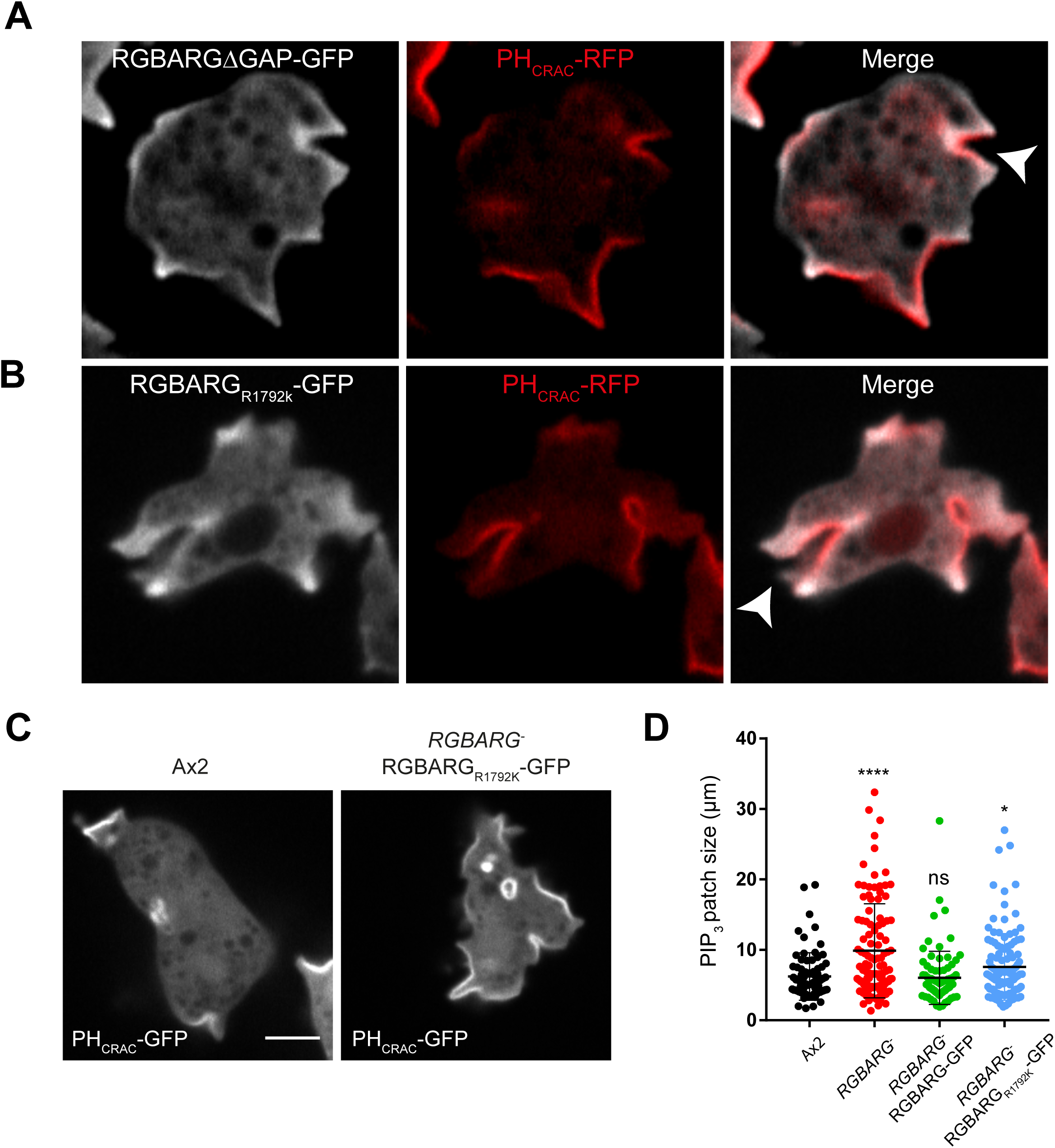
(A) Co-expression of RGBARGdRasGAP-GFP and PHCRAC-RFP in RGBARG-cells indicating that the RasGAP domain helps excluded RGBARG-GFP from PIP3 rich regions of the cell. (B) The GAP activity inactivating point mutant R1792K localizes normally and is excluded from the base of protruding cups (arrowhead). (C) PHCRAC-GFP localization in Ax2 and RGBARG-cells expressing RGBARGR1792K-GFP, demonstrating that PIP3 dynamics are not rescued by this construct. PHCRAC-GFP patch size is quantified in (D).

**Figure 5 Supplement:**
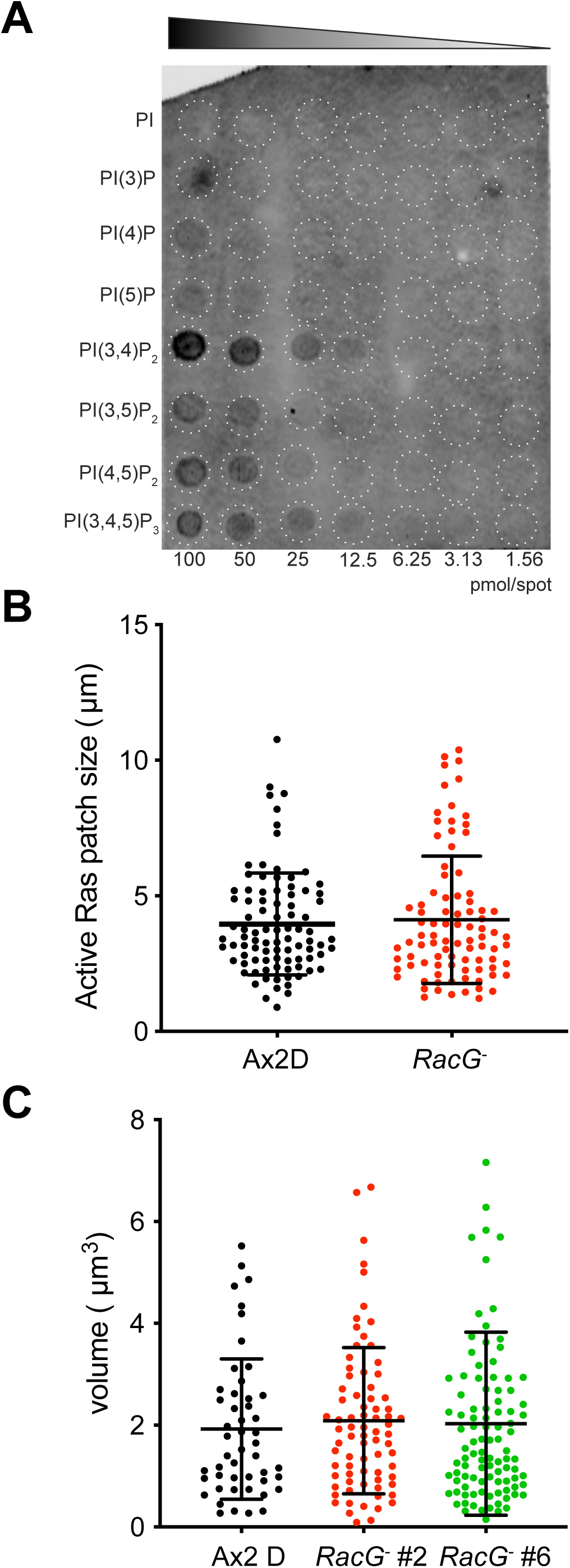
(A) PIP array analysis of BAR-GFP binding showing a moderate preference for PI(3,4)P2 in this assay. (B) Active Ras signaling during macropinosome formation in RacG-cells and their parental cell line Ax2D. Patch size was quantified from single confocal planes of cells expressing RBD-GFP. (C) Volume of macropino-somes formed by RacG- and control cells. Measured by imaging cells taking up FITC dextran. Two independent RacG-clones were analysed over 3 independent experiments.

## Notes

#### Summary of Updates

Rectification of the omission of David Traynor as co-author, and inclusion of an additional model figure to summarise our findings.

